# A Glutamate-Neuropeptide Circuit Converts Transient Mate Contact into a Persistent Local Search State in *Caenorhabditis elegans* Males

**DOI:** 10.64898/2026.11.10.762718

**Authors:** Yu-Han Chen, Yin-Chen Lin, Tse-Yu Chen, Kai-An You, Wei-En Ma, Ying-Syuan Lai, Chieh Wang, Jia-Bin Yang, Chun-Hao Chen

## Abstract

Transient sensory encounters can trigger behavioral states that persist long after the initiating stimulus has disappeared, yet how neural circuits initiate and sustain these states remains poorly understood. Here, we show that brief contact with a suitable mate induces a persistent local search state in male *Caenorhabditis elegans*, characterized by increased reversals and spatially restricted exploration that continues after mate removal. Transient optogenetic activation of male-specific ray type B (RnB) sensory neurons recapitulates this state, with its magnitude and persistence increasing with stimulation strength. Glutamatergic signaling promotes neuronal responses within the interconnected RnB sensory network and contributes to prolonged activity in downstream LUA interneurons. In contrast, loss of the LUA-derived neuropeptide NLP-13 or its receptor DMSR-2 preserves initial stimulus-evoked neuronal responses but reduces prolonged LUA calcium activity and behavioral persistence. LUA-specific restoration of NLP-13 rescues persistent local search, identifying these interneurons as a functionally important source of the neuropeptide. Together, our findings reveal complementary contributions of glutamatergic and neuropeptidergic signaling to the generation and persistence of behavioral states, illustrating how distinct signaling mechanisms cooperate to transform transient social encounters into prolonged circuit activity and persistent behavior.

## Introduction

Animals transform transient sensory experiences into behavioral states that persist beyond the initiating stimulus, allowing recent encounters to influence subsequent actions over extended periods ^1, 2, 3, 4, 5^. Generating such states involves two fundamental processes: integrating sensory information to initiate a behavioral transition and maintaining the resulting state after sensory input has ceased. Persistent behavioral states have been associated with prolonged neuronal activity, recurrent circuit dynamics, and neuromodulatory signaling ^2, 3, 6, 7, 8, 9, 10^. However, how these mechanisms coordinate to initiate and sustain behavioral states, and whether distinct signaling pathways govern these processes, remains poorly understood.

Studies across animal systems have revealed neural circuits that convert transient sensory input into persistent behavioral responses. In male *Drosophila*, activation of P1 neurons by female-associated sensory cues induces persistent arousal, while downstream pCd neurons contribute to sustained courtship- and aggression-related behaviors ^3, 6^. In larval zebrafish, persistent neural dynamics distinguish exploratory and exploitative phases of foraging, with dorsal raphe neurons exhibiting activity associated with exploitation ^9^. These findings highlight the importance of sensory integration and prolonged circuit activity in generating behavioral persistence. At the molecular level, fast synaptic transmission can rapidly recruit interconnected neurons, whereas neuromodulators, including neuropeptides, can influence circuit excitability over longer timescales ^8, 11, 12, 13, 14, 15, 16^. In *Caenorhabditis elegans*, serotonin and pigment-dispersing factor signaling regulate persistent roaming and dwelling states, while interactions between multisensory inputs and neuropeptide signaling organize search-related behaviors ^8, 17^. Despite these advances, how fast neurotransmission and neuropeptide signaling cooperate to transform transient sensory input into persistent circuit activity and sustained behavioral responses remains elusive.

The compact and well-characterized nervous system of *C. elegans* provides an opportunity to dissect these mechanisms at cellular and molecular resolution ^18, 19^. In addition to the neurons shared with hermaphrodites, males possess sex-specific sensory circuits specialized for reproductive behaviors ^18, 19, 20^. Male *C. elegans* actively search for mating partners and rely on specialized tail sensory neurons to detect contact-dependent cues and initiate mating responses ^21, 22, 23, 24, 25, 26^. We previously demonstrated that mate recognition involves a two-step sensory process in which males detect cuticular surface cues and subsequently evaluate the mechanical stiffness of the contacted animal ^21^. Such contact-dependent recognition provides a biologically relevant context in which encounters with potential mates could influence subsequent exploratory behavior. However, it is unclear how mate contact induces a behavioral state that persists after physical separation, and how the underlying sensory circuits generate and maintain such persistence.

Here, we show that brief contact with a suitable mate induces a persistent local search state in male *C. elegans*, characterized by increased reversals and spatially restricted exploration that continues after mate removal. Transient activation of male-specific ray type B (RnB) sensory neurons is sufficient to induce this state, with its magnitude and persistence increasing with stimulation strength. Glutamatergic signaling facilitates neuronal responses within the RnB sensory network and contributes to prolonged activity in downstream LUA interneurons. In contrast, the LUA-derived neuropeptide NLP-13 and its receptor DMSR-2 support prolonged LUA calcium activity and behavioral persistence. Together, our findings reveal complementary glutamatergic and neuropeptidergic mechanisms that contribute to the generation and persistence of mate-induced local search, providing a circuit-level framework for understanding how transient sensory experiences generate lasting behavioral responses.

## Results

### Brief mate contact induces a persistent local search state

To study neural mechanisms of contact-mediated mate recognition, we positioned mates in close proximity to moving males and allowed a 30-s contact period for mate recognition. Then, we removed mates and observed behavioral changes in males (Fig. 1a). Before contact, males primarily displayed forward locomotion with intermittent pauses and reversals. Encounters with an immobilized *unc-13* mutant hermaphrodite, compared with mock treatment (no contact), increased reversal frequency without significantly altering turning frequency (Fig. 1b, c; Supplementary Fig. 1a). These behavioral responses resulted in more spatially restricted trajectories (Fig. 1b, Supplementary Video 1). Instantaneous speed was not significantly altered by contact, suggesting a change in locomotor patterns rather than overall movement speed (Supplementary Fig. 1b). In addition, unsupervised behavioral classification by CASTLE further revealed increased occupancy of curling and looping motifs during the post-contact period (Fig. 1d; Supplementary Fig. 1c), capturing behavioral features not detected by our conventional locomotion analyses ^27^. Because we operationally defined successful contact as 30 s of continuous interaction, males could make brief contacts and initiate behavioral changes before reaching this criterion, leading to a gradual rather than sharply demarcated transition around the defined contact period. Together, these changes produced a sustained reduction in spatial exploration that persisted for several minutes after mate removal and gradually returned to exploratory locomotion (Fig. 1e-g). We refer to this persistent and spatially restricted locomotor program as the mate-induced local search state. To quantify overall behavioral changes, we calculated the ratio of net displacement in each 10-s bin relative to the individual pre-stimulation mean as a locomotion index (Fig. 1e, h). Consistent with our behavioral characterization, the locomotion index decreased significantly upon contact with an immobilized *unc-13* mutant hermaphrodite as the potential mate (Fig. 1h; Supplementary Video 1). Overall, these data suggest that males enter a persistent local-search behavioral state characterized by restricted spatial exploration after mate contact.

**Fig. 1.**
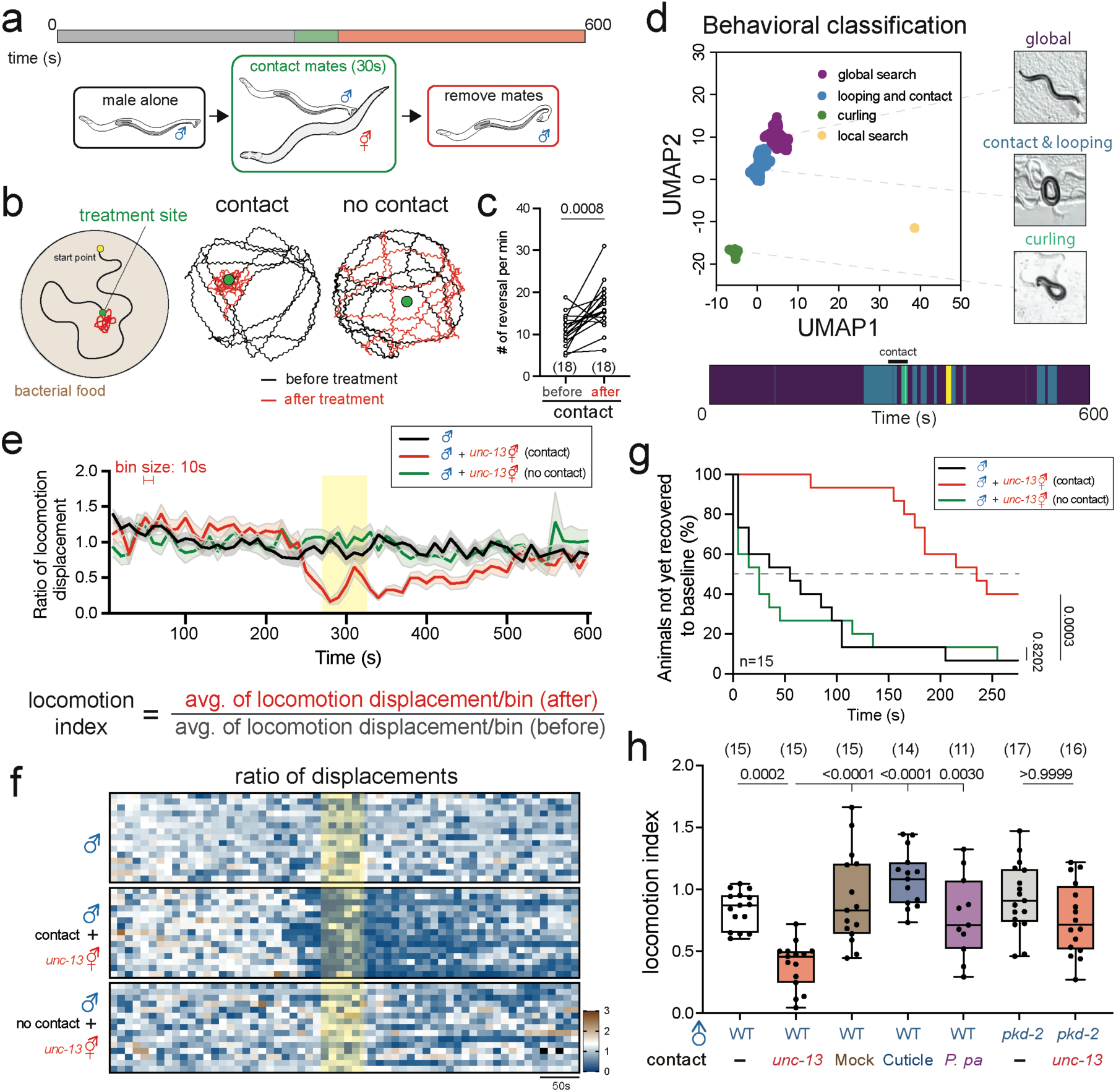
Contact with a mate induces a persistent local search state in *C. elegans* males. a, Schematic of the contact assay. A freely moving male was recorded alone, allowed to contact a mate for 30 s, and subsequently recorded after removal of the mate. b, Representative locomotion trajectories before (black) and after (red) contact with a mate. The green circle indicates the treatment site and the yellow circle indicates the starting position. c, Number of reversals per minute before and after contact with a mate. Lines connect measurements from the same animal. Statistical significance was assessed using a two-tailed paired Student’s *t*-test. d, Unsupervised behavioral classification of male locomotion. UMAP representation separates global search, looping/contact, curling and local-search behaviors (top left), with representative postures shown at right. The behavioral-state sequence of a representative animal over the 600-s recording period is shown below. e, Time course of the locomotion displacement ratio for males tested alone (black), males following contact with *unc-13* hermaphrodites (red) and males exposed to *unc-13* hermaphrodites without contact (green). Locomotion was quantified in 10-s bins. The yellow region denotes the 30-s treatment period. The locomotion displacement ratio was calculated as the displacement in each 10-s bin/individual pre-stimulation mean. Lines indicate the mean and shaded regions indicate variability (±SEM) among animals. f, Heatmaps showing the temporal dynamics of locomotion for individual males under the conditions in e. The yellow region indicates the treatment period. g, Time to baseline displacement recovery following mate removal. Curves show the percentage of animals that had not yet met the baseline-recovery criterion for males tested alone (black), males following contact with *unc-13* hermaphrodites (red) and males exposed to *unc-13* hermaphrodites without contact (green); *n =* 15 per condition. The dashed line indicates 50% not yet recovered. Kaplan–Meier recovery curves were compared using the log-rank (Mantel–Cox) test. h, Locomotion indices following the indicated contact and control treatments, including mock treatment, cuticle exposure, *P. pacificus* exposure and experiments using *pkd-2* and *unc-13*. Boxes indicate the median and 25th-75th percentiles, whiskers indicate the minimum and maximum values, and dots represent individual animals. Statistical significance was assessed by one-way ANOVA with Bonferroni correction for multiple comparisons. Sample sizes are indicated above each group. Exact *P* values are shown in the figure.

In *C. elegans*, males are attracted by secreted ascaroside and volatile pheromones through modulating their locomotion behaviors ^28, 29, 30^. Consequently, we considered whether residual pheromones at the contact site triggered the transition between behavioral states. To test this, we positioned an *unc-13* immobilized mutant hermaphrodite away from the male to prevent physical contact and found that the male did not display the local search state (Fig. 1e, h). In contrast, brief contacts of the *daf-22* mutant hermaphrodite, defective in ascaroside pheromone production ^31^, still evoked the local search state of males (Supplementary Fig. 2). The behavioral switch was also observed in males with mutations in *srd-1* (Supplementary Fig. 2), a receptor for volatile pheromones ^30^, indicating that ascarosides and SRD-1-dependent volatile pheromone signaling are dispensable for contact-induced local search.

In our previous study, we showed that males use a two-step sensory mechanism involving cuticular cues and body stiffness to identify compatible mates via physical contact ^21^. Consistent with this view, males did not transition to local search behaviors after contacting either purified cuticles or a non-recognizable nematode species, *Pristionchus pacificus*, which possesses comparable body stiffness (Fig. 1h) ^21^. These observations are consistent with our previous finding that recognition requires integrating cuticular and mechanical information. Together, these results suggest that entry into the local search state depends on recognizing a suitable mate rather than on physical contact or pheromonal cues alone. We therefore asked which contact-sensing neurons are sufficient to convert this brief sensory encounter into a persistent state.

### Transient RnB activation is sufficient and promotes entry into the local search state

Because males physically contact hermaphrodites to access these sensory cues, we asked whether direct neuronal activation could bypass physical contact and induce a persistent behavioral state. Males sense the physical presence of hermaphrodites mainly through 9 pairs of tail rays. Each ray comprises A- and B-type ray sensory neurons ^32^. PKD-2 is the homolog of polycystin-2 expressed in male tails that mediates the contact responses ^33^. Confirming our hypothesis, contact-defective *pkd-2* mutant males were unable to transition into the local search state (Fig. 1h), suggesting that ray neurons are involved ^33^. To further study whether transient activation of ray sensory neurons induces the same behavioral persistence as brief contact of hermaphrodites, we expressed a red-light-sensitive cation channel Chrimson under the control of the *tba-9* and the *pkd-2* promoters to activate RnAs and RnBs, respectively ^34^. A 500-ms red-light activation of RnAs and RnBs resulted in characteristic curls of male tails as previously reported ^26^, validating the efficacy of Chrimson-mediated activation of ray neurons (Supplementary Video 2 and 3). Initially, individual males were allowed to move freely and we then transiently activated RnA or RnB neurons optogenetically using various numbers of pulses (Fig. 2a). We found that 20 pulses of RnA activation (500-ms stimulation separated by a 1-s pause) did not lead to a significant alteration in the locomotion index and dynamics compared to the mock control without the co-factor all-trans-retinal (ATR) (Supplementary Fig. 3). Conversely, employing the same activation protocol for RnB neurons resulted in a marked reduction in locomotion displacements (Fig. 2b, c). Characterization of behavioral changes triggered by RnB activations also revealed similar changes in turning and reversal frequencies, together with increased occupancy of looping and curling behavioral classes (Fig. 2d; Supplementary Fig. 4, Supplementary Video 4). The RnB-induced reduction in locomotor displacement was maintained for several minutes, resembling the behavioral response following brief contact with hermaphrodites (Fig. 2e).

**Fig. 2.**
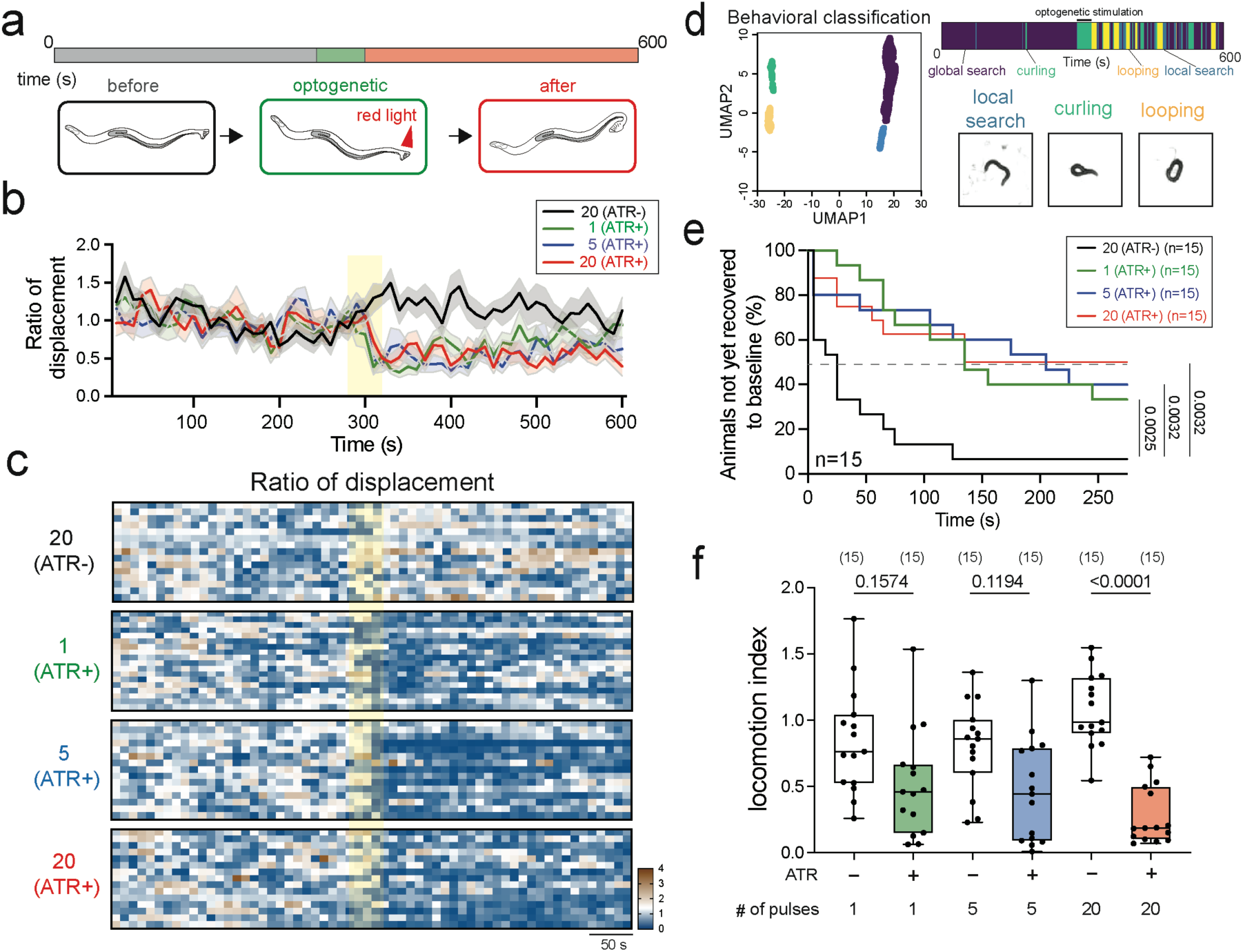
Optogenetic activation of male sensory neurons is sufficient to induce a persistent local search state. a, Schematic of the optogenetic stimulation assay. Male locomotion was recorded before, during and after red-light stimulation. b, Time course of the locomotion displacement ratio following 1, 5 or 20 optogenetic stimulations in ATR-positive animals and following 20 stimulations in ATR-negative controls. The yellow region denotes the stimulation period. Lines indicate the mean and shaded regions indicate variability (±SEM) among animals. c, Heatmaps showing locomotion dynamics of individual animals following the indicated stimulation protocols. d, UMAP-based behavioral classification following optogenetic stimulation, identifying global search, curling, looping and local-search states. Representative animal postures are shown at right, and the behavioral-state sequence over the 600-s assay is shown above. e, Time to baseline displacement recovery following 1, 5 or 20 stimulations in ATR-positive animals compared with 20-stimulation ATR-negative controls. Curves show the percentage of animals that had not yet met the baseline-recovery criterion; *n =* 15 per condition. The dashed line indicates 50% not yet recovered. Kaplan–Meier recovery curves were compared using the log-rank (Mantel-Cox) test. f, Locomotion indices following paired stimulation protocols in ATR-negative and ATR-positive animals. Boxes indicate the median and 25th-75th percentiles, whiskers indicate the minimum and maximum values, and dots represent individual animals. Statistical significance was assessed by two-way ANOVA with Bonferroni correction for multiple comparisons. Sample sizes are indicated above the plots. Exact *P* values are shown in the figure.

Persistent behavioral states often vary in magnitude or duration with the strength of the initiating stimulus. We therefore asked whether local search depends on the extent of RnB activation. Activation of RnBs with 1- or 5-pulse stimuli produced only modest reductions in locomotion index, whereas 20-pulse stimulation induced a substantially stronger and more persistent response (Fig. 2b, c, e, f). Thus, RnB activity is sufficient to induce the local search state, with its magnitude and persistence increasing with repeated RnB activation. Together with the inability of RnA activation alone to induce a comparable state, these findings identify RnBs as the principal ray sensory population sufficient to drive entry into persistent local search.

### Glutamatergic transmission contributes to RnB-induced local search

Ray neurons form an extensively interconnected network through both chemical synapses and gap junctions (Fig. 3a) ^18^. To determine whether chemical synaptic transmission from RnB neurons is required for local search, we expressed tetanus toxin light chain in RnBs to inhibit synaptic vesicle release (Fig. 3b). Disrupting neurotransmission from RnBs nearly abolished the local search state, indicating that chemical synaptic output from RnBs is critical for recruiting downstream circuits. Previous studies have shown that RnB neurons collectively express a diverse repertoire of neurotransmitters, including acetylcholine, γ-aminobutyric acid (GABA), glutamate, serotonin, and octopamine (Fig. 3c) ^35^. To delineate the neural circuit downstream of RnB neurons, we first interrogated the role of major neurotransmitters by optogenetically activating RnBs in mutants defective in glutamate, serotonin, or octopamine. Acetylcholine and GABA were excluded because these mutants exhibit severe locomotor defects that would confound our behavioral analysis. Notably, the absence of *eat-4*, which encodes the vesicular glutamate transporter, prevented males from transitioning into the local search state (Fig. 3d; Supplementary Video 5) ^36^. We confirmed the functional requirement of *eat-4* by restoring its activity using a 5.5 kb upstream promoter, which rescued the behavioral defects (Fig. 3d). To identify the site of glutamatergic action, we used cell-specific RNAi to selectively knock down *eat-4* in RnB or its downstream glutamatergic LUA interneurons. Knockdown of *eat-4* in RnBs, but not LUAs, significantly impaired the transition into the persistent behavioral state (Fig. 3d), supporting a contribution of RnB-derived glutamatergic signaling to the behavioral response. However, restoring *eat-4* expression in RnBs, LUAs, or both neuronal populations failed to rescue the *eat-4* mutant phenotype (Fig. 3d). These findings suggest that glutamatergic signaling from RnBs contributes to the establishment of the local search response, while the failure of cell-specific rescue raises the possibility that additional glutamatergic neurons or broader *eat-4* expression are required for the full behavioral response.

**Fig. 3.**
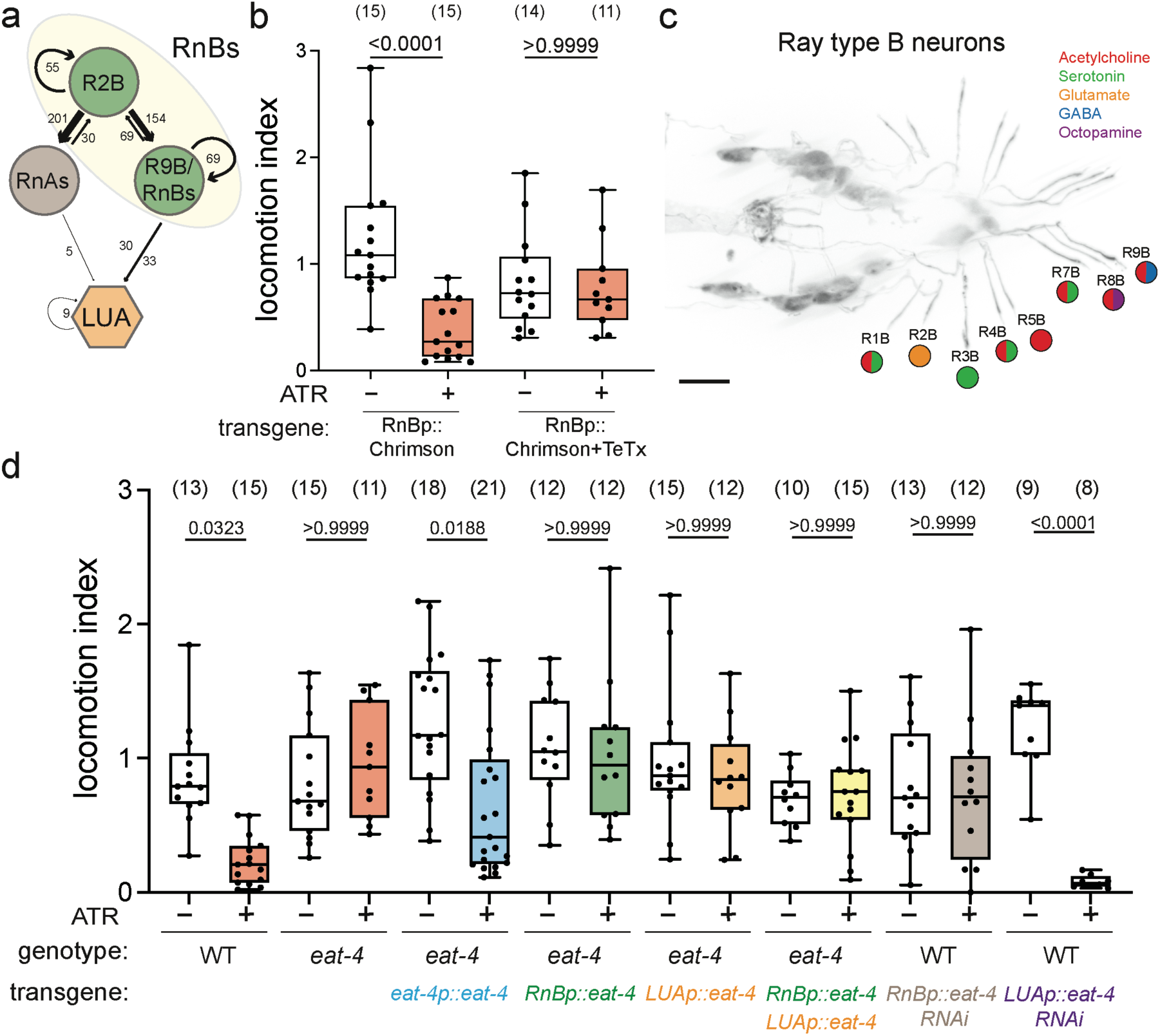
Glutamatergic transmission contributes to RnB-induced persistent local search. a, Schematic of connectivity among R2B, the ray type A neurons (RnAs), ray type B neurons (RnBs) and the LUA interneurons. Numbers indicate annotated synaptic connections between the indicated neuronal classes. b, Locomotion indices following optogenetic stimulation in control males and males expressing tetanus toxin (TeTx) in RnB neurons. ATR-negative and ATR-positive animals are shown for each genotype. c, Neurotransmitter identities of individual ray type B neurons. Pie-chart colors indicate reported cholinergic, serotonergic, glutamatergic, GABAergic and octopaminergic identities. Scale bar = 10 μm. d, Locomotion indices following optogenetic stimulation in wild-type and *eat-4* mutant males and following tissue-specific restoration or knockdown of *eat-4*. Conditions include pan-*eat-4* rescue, RnB-specific rescue, LUA-specific rescue, combined RnB- and LUA-specific rescue and tissue-specific RNAi, as indicated beneath the plot. Boxes indicate the median and 25th-75th percentiles, whiskers indicate the minimum and maximum values, and dots represent individual animals. Statistical significance was assessed by two-way ANOVA with Bonferroni correction for multiple comparisons. Sample sizes are indicated above individual groups and exact *P* values are shown in the figure.

In addition to glutamate signaling, we found that serotonin-deficient *tph-1* mutants still transitioned into the local search state following RnB stimulation, whereas octopamine-deficient *tbh-1* mutants exhibited reduced RnB-evoked local search (Supplementary Fig. 5), indicating that octopaminergic signaling further modulates this behavioral state ^35, 37^. Because the relevant octopaminergic circuitry remains unresolved, we focused subsequent analyses on the glutamatergic pathway linking RnBs to LUA.

### LUA activity is required for and modulates RnB-induced local search

We next asked which downstream neurons integrate RnB sensory output to control entry into the local search state. Given that LUA interneurons are the immediate synaptic targets of RnBs, express glutamate receptors, and have been implicated in persistent search behaviors (Fig. 3a) ^18, 38, 39^, we examined their role in this circuit. To determine whether LUA activity is necessary for the behavioral state, we silenced LUA neurons using a histamine-gated chloride channel (HisCl1) ^40^. Application of histamine efficiently abolished the behavioral switch upon RnB activation, whereas transgenic animals behaved comparably to wild-type controls in the absence of histamine (Fig. 4a). These data indicate that LUA function is required for RnB-induced persistent local search. However, optogenetic activation of LUAs alone was insufficient to induce the local search state (Fig. 4b), suggesting that LUAs function as a critical circuit node rather than acting alone to command the behavioral transition. Consistent with a role for LUA excitability in regulating state entry, sustained enhancement of LUA activity through expression of *pkc-1(gf)* sensitized males to the behavioral response following a 5-pulse stimulus paradigm (Fig. 4c) ^41^. Together, these findings identify LUA as a downstream node whose excitability modulates responsiveness to RnB input.

**Fig. 4.**
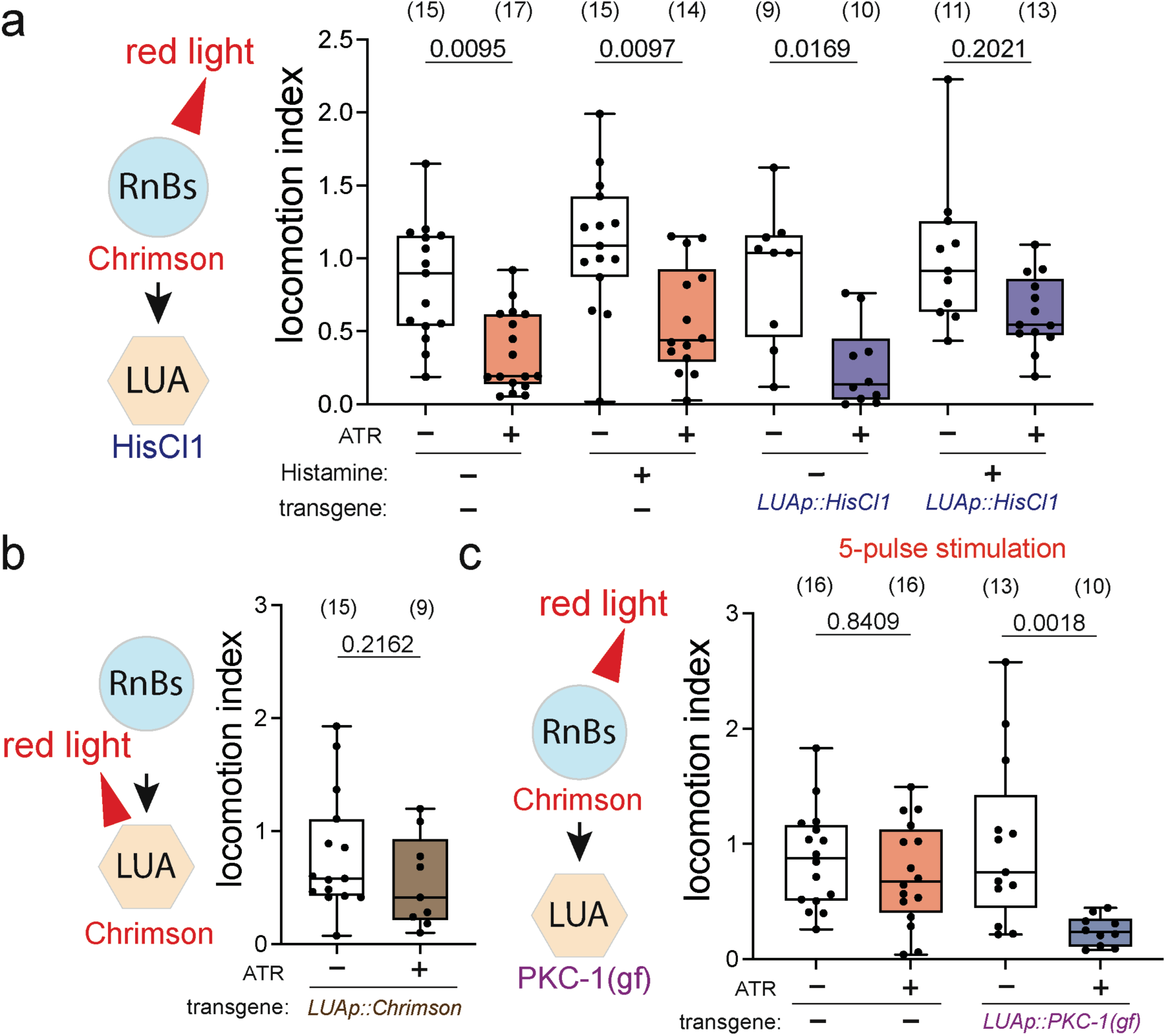
LUA interneurons are required for, but are not by themselves sufficient to initiate, the persistent behavioral state. a, Left, experimental schematic for optogenetic activation of RnB neurons while inhibiting LUA neurons using HisCl1. Right, locomotion indices of ATR-negative and ATR-positive animals in the absence or presence of histamine, with or without LUA-specific HisCl1 expression. b, Left, schematic of direct optogenetic activation of LUA neurons. Right, locomotion indices of ATR-negative and ATR-positive males expressing Chrimson in LUA. c, Left, experimental schematic for optogenetic activation of RnB neurons combined with expression of constitutively active PKC-1, PKC-1(gf), in LUA. Right, locomotion indices following five optogenetic stimuli in control animals and males expressing LUA-specific PKC-1(gf). Boxes indicate the median and 25th-75th percentiles, whiskers indicate the minimum and maximum values, and dots represent individual animals. Statistical significance was assessed by two-way ANOVA with Bonferroni correction for multiple comparisons (a, c) or a two-tailed unpaired Student’s t test (b). Sample sizes are indicated above each group and exact *P* values are shown in the figure.

### Glutamatergic signaling modulates RnB and LUA calcium responses

We next investigated how RnB sensory neurons relay information to LUA interneurons to sustain persistent circuit activity by calcium imaging of GCaMP8f ^42^. Because fluorescence intensity gradually decayed over the recording period, we evaluated approaches to correct for photobleaching. However, extrapolation-based correction of the pre-stimulation decay systematically introduced an artificial upward shift in the post-stimulation signal. We therefore present the uncorrected ΔF/F_0 t_races and quantified stimulus-evoked and persistent responses using predefined within-recording analysis windows. Optogenetic activation of RnBs elicited a clear GCaMP8f response in LUA neurons during the stimulation window, consistent with the functional coupling between these sensory neurons and downstream interneurons (Fig. 5a; Supplementary Fig. 6a, Supplementary Fig. 7). Compared with the ATR-negative control, RnB stimulation in wild-type males produced a prolonged calcium-associated fluorescence response that remained relatively elevated during the post-stimulation period (120-240 s; Fig. 5a, b; Supplementary Fig. 6a). These observations suggest that transient RnB activation induces prolonged changes in LUA calcium dynamics associated with the persistent local search state.

**Fig. 5.**
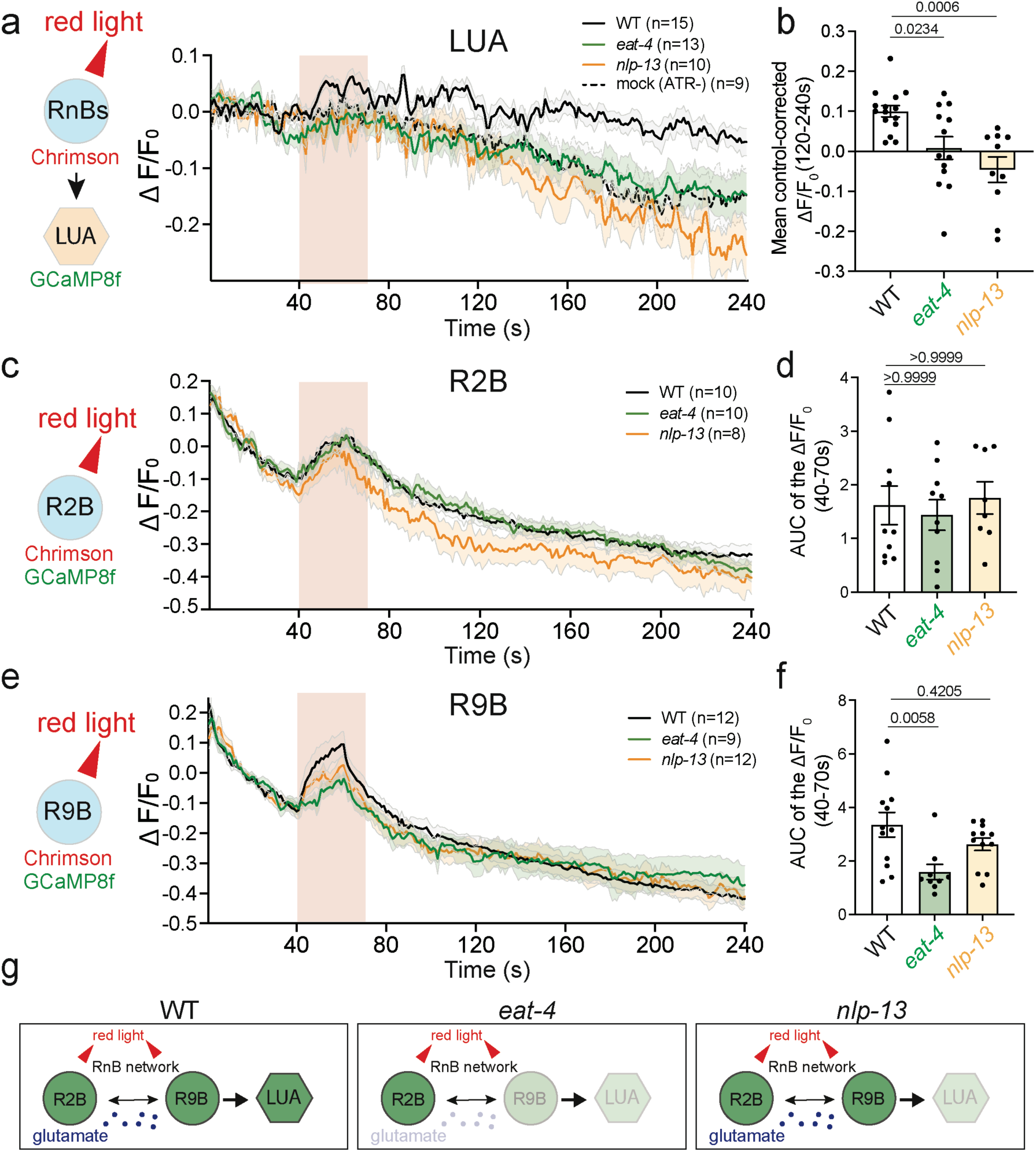
Glutamate and NLP-13 differentially regulate sensory-network activation and persistent LUA activity. a, Calcium responses of LUA neurons during optogenetic activation of RnB neurons in wild-type, *eat-4* and *nlp-13* males. Animals without all-trans-retinal (ATR-) served as unstimulated controls (mock). Traces show ΔF/F_0_ over time; the shaded red region denotes the stimulation period. Lines indicate mean responses and surrounding shading indicates variability among animals. b, Mean control-corrected LUA ΔF/F_0_ during the late post-stimulation period (120-240 s). At each time point, the mean ΔF/F_0_ of ATR-negative controls was subtracted from the corresponding fluorescence value of each animal. Corrected values were then averaged across the indicated interval to obtain one value per animal. Bars represent mean ±SEM, and dots indicate individual animals. Statistical significance was assessed using one-way ANOVA with Bonferroni correction for multiple comparisons. Exact P values are shown. c, Calcium responses of R2B during direct optogenetic activation of the RnBs population in wild-type, *eat-4* and *nlp-13* males. d, Area under the curve (AUC) of R2B calcium responses during the stimulation-associated interval (40-70 s). e, Calcium responses of R9B during optogenetic stimulation of the RnBs population in wild-type, *eat-4* and *nlp-13* males. f, AUC of R9B calcium responses during the stimulation-associated interval (40-70 s). g, Summary of optogenetically evoked calcium responses in RnB sensory neurons and downstream LUA interneurons in wild-type, *eat-4*, and *nlp-13* males. Data are mean ±SEM with individual animals shown. Statistical significance was assessed by one-way ANOVA with Bonferroni correction for multiple comparisons. Sample sizes are indicated in the corresponding traces, individual data points are shown in the summary plots and exact *P* values are indicated.

Among the nine bilateral pairs of RnB neurons, R2B is the only pair reported to use glutamate as a neurotransmitter ^35^. Notably, R2B does not directly synapse onto LUA but is extensively connected to other Ray type B neurons, positioning it as a potential source of glutamatergic signaling within the RnB network ^18, 43^. In contrast, R9B forms the strongest anatomical connection with LUA among the RnB neurons (Supplementary Fig. 6b) ^18, 43^. We therefore optogenetically activated RnBs in *eat-4* mutants and monitored calcium responses in R2B and R9B as representative nodes along the proposed sensory-network relay. This comparison allowed us to distinguish activation of the putative glutamatergic source neuron itself from glutamate-dependent recruitment of an RnB neuron that is strongly connected to the downstream LUA interneurons. R2B calcium responses were comparable between wild-type and *eat-4* mutant males (Fig. 5c, d; Supplementary Fig. 6a), indicating that glutamatergic transmission is not required for the initial activation of R2B itself. By contrast, the stimulus-evoked calcium response in R9B was reduced in *eat-4* mutants, consistent with a model in which glutamatergic signaling enhances the responsiveness of other RnB neurons (Fig. 5e, f, g; Supplementary Fig. 6a). Although *eat-4* mutants retained a comparable acute LUA calcium response during the direct optogenetic stimulation window (40-70 s) (Supplementary Fig. 7), *eat-4* mutants exhibited a significantly reduced late LUA fluorescence response relative to the ATR-negative control, consistent with impaired prolonged circuit activation following RnB stimulation (Fig. 5a, b; Supplementary Fig. 7). Consistent with this, *eat-4* mutants exhibited little sustained reduction in locomotor displacement and rapidly returned to baseline following RnB stimulation, reflecting impaired establishment of the local search state (Fig. 6a, b). Together, these results suggest that glutamatergic signaling facilitates neuronal responses within the RnB sensory network and supports prolonged activity in downstream LUA interneurons (Fig. 5g), thereby contributing to the generation of the persistent local search state.

**Fig. 6.**
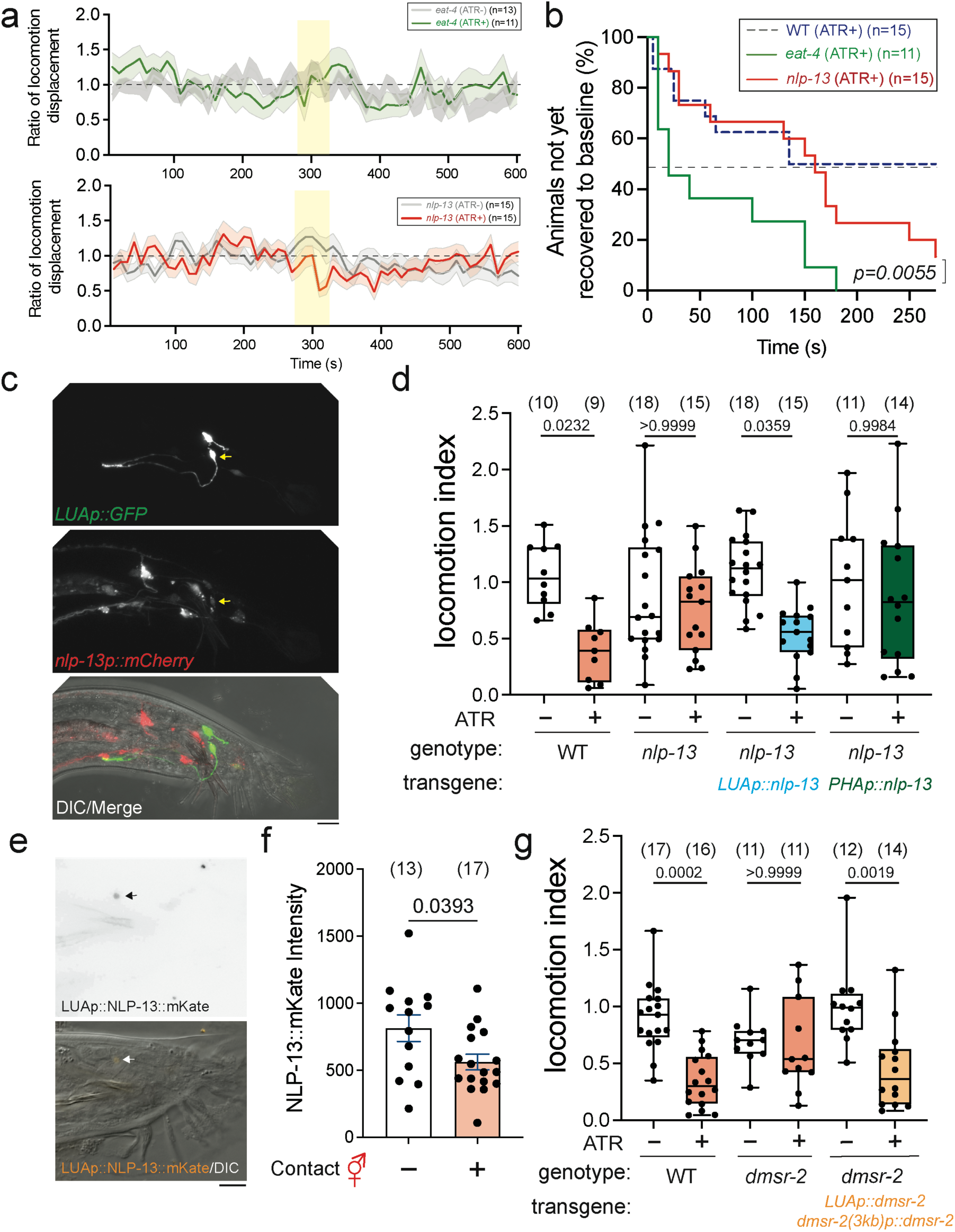
**NLP-13 neuropeptide signaling promotes persistence of the mate-induced behavioral state**. a, Time courses of locomotion displacement ratios in *eat-4* (top) and *nlp-13* (bottom) males in ATR-negative and ATR-positive conditions. The yellow region denotes the stimulation period. Lines indicate mean responses and shaded regions indicate variability (±SEM) among animals. b, Time to baseline displacement recovery in optogenetically stimulated wild-type (adapted from Fig. 2e), *eat-4* and *nlp-13* males. Curves indicate the percentage of animals that had not yet met the baseline-recovery criterion; the dashed line indicates 50% not yet recovered. Kaplan–Meier recovery curves were compared using the log-rank (Mantel-Cox) test. c, Representative images showing LUA-specific GFP (top), *nlp-13p::mCherry* (middle) and merged DIC/fluorescence signals (bottom). Arrows indicate the LUA region. Scale bar = 10 μm. d, Locomotion indices in wild-type and *nlp-13* mutant males and following LUA-specific or PHA-specific expression of *nlp-13*. ATR-negative and ATR-positive conditions are shown. e, f, NLP-13::mKate fluorescence was measured in males with or without 2-h contact with hermaphrodites (e, representative fluorescence and DIC images; f, quantification). NLP-13::mKate fluorescence was reduced following contact. Scale bar = 10 μm. Data are mean ±SEM with individual animals shown. Statistical significance was assessed using a two-tailed unpaired Student’s *t*-test. g, DMSR-2 regulates the persistent local search state. Locomotion indices of wild-type and *dmsr-2* mutant males, together with *dmsr-2* mutants expressing the receptor under the control of a 3-kb upstream regulatory region and a LUA-specific promoter. Combined expression rescued the *dmsr-2* mutant phenotype. Boxes indicate the median and interquartile range, whiskers indicate the minimum and maximum values, and dots represent individual animals. Statistical significance was assessed using two-way ANOVA with Bonferroni correction for multiple comparisons. Sample sizes and exact *P* values are indicated in the figure.

### LUA-derived NLP-13 neuropeptide signaling sustains the local search state

We next asked how LUA neurons maintain persistent activity after the initial sensory stimulus. Although LUA neurons are glutamatergic, LUA-specific knockdown of *eat-4* did not attenuate the local search state (Fig. 3d), suggesting that LUA-derived glutamatergic signaling is not essential for the behavioral response under these conditions. Because neuropeptide signaling can sustain neuronal activity over prolonged timescales, we hypothesized that peptidergic signaling from LUA contributes to maintenance of the local search state. We therefore focused on *nlp-13*, which encodes a neuropeptide-like protein enriched in LUA neurons (Fig. 6c) ^44^. Loss of *nlp-13* significantly reduced the change in locomotor displacements associated with the local search state (Fig. 6d; Supplementary Video 6). Restoring *nlp-13* expression specifically in LUA neurons fully rescued this defect, but not in neighboring PHA neurons. These data indicate that LUA is a functionally relevant source of NLP-13. Consistent with this model, NLP-13::mKate (translational fused peptide) fluorescence decreased in LUA neurons after 2 h of prolonged contact with hermaphrodites, suggesting stimulus-dependent depletion or redistribution of the NLP-13 pool in LUA (Fig. 6e, f). Because PHA neurons lie in close proximity to LUA, the failure of PHA-derived NLP-13 to restore the behavioral response also suggests that local peptide expression alone is insufficient. Instead, effective NLP-13 signaling depends on the appropriate cellular source for peptidergic transmission. We thus conclude that NLP-13 neuropeptide signaling from LUA neurons supports persistent local search.

### NLP-13-DMSR-2 signaling supports prolonged LUA activity

To further characterize the contribution of NLP-13 to behavioral persistence, we examined the temporal dynamics of locomotor recovery following RnB stimulation (Fig. 6a). In *eat-4* mutants, RnB stimulation elicited little sustained reduction in locomotor displacement, and animals rapidly returned to their baseline displacement (median recovery time, 20 s). In contrast, *nlp-13* mutants exhibited an initial reduction in locomotor displacement followed by gradual recovery (median recovery time, approximately 150 s; Fig. 6a, b). Because *eat-4* mutants exhibited little initial local search response, their rapid return to baseline likely reflects impaired establishment of the local search state rather than accelerated termination of an established state. In contrast, *nlp-13* mutants displayed an initial behavioral response followed by an accelerated return toward baseline compared with wild-type, consistent with impaired behavioral persistence. These distinct profiles suggest that glutamatergic and NLP-13 signaling make differential, although potentially overlapping, contributions to the establishment and maintenance of persistent local search.

Calcium imaging further revealed that *nlp-13* mutants retained stimulus-evoked responses in R2B and R9B but exhibited reduced prolonged LUA calcium activity (Fig. 5; Supplementary Fig. 6 and 7). Together, these findings indicate that NLP-13 signaling contributes to prolonged circuit activity and behavioral persistence but is not essential for the initial stimulus-evoked neuronal responses. Although these observations are consistent with differential contributions of glutamatergic and neuropeptidergic signaling to state establishment and persistence, they do not establish that the two pathways act exclusively during distinct phases of the behavioral response.

**Fig. 7.**
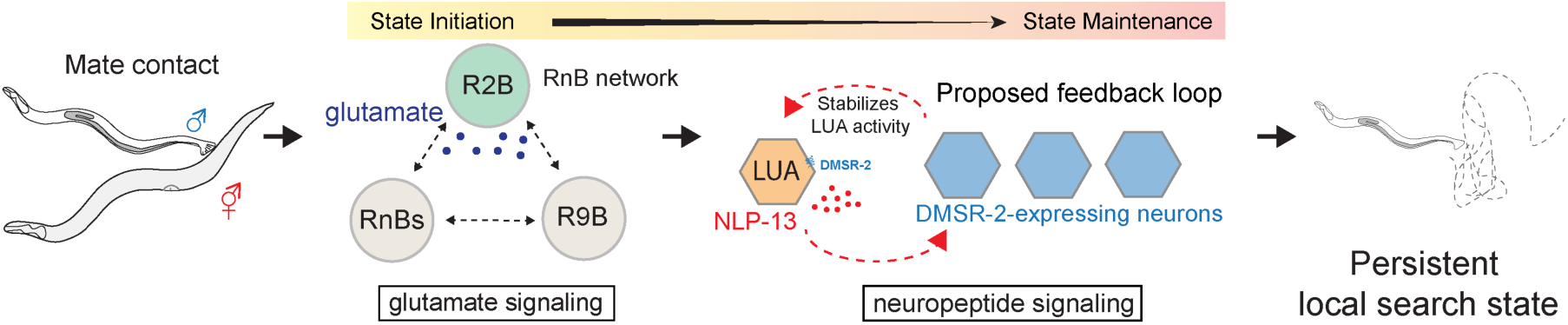
Model for the glutamate-neuropeptide circuit that converts transient mate contact into a persistent local search state. Contact with a suitable mate activates the male ray type B (RnB) sensory network. Glutamatergic signaling contributes to neuronal responses within the interconnected RnB network, including R2B and R9B, and supports prolonged activity in downstream LUA interneurons. LUA-derived NLP-13 engages DMSR-2-dependent signaling that contributes to prolonged LUA activity and behavioral persistence. These glutamatergic and neuropeptidergic mechanisms may operate across overlapping temporal phases to support the generation and persistence of the local search state following transient mate contact.

Having established LUA-derived NLP-13 as a contributor to behavioral persistence, we next investigated the receptor mediating this peptidergic signal. DMSR-2 has previously been identified as a G protein-coupled receptor for NLP-13 ^45^. Consistent with this relationship, *dmsr-2* mutants exhibited impaired local search resembling the phenotype of *nlp-13* mutants (Fig. 6g). Expression of *dmsr-2* driven by either a 3-kb endogenous upstream regulatory region or a LUA-specific promoter alone failed to restore behavioral persistence (Supplementary Fig. 8). In contrast, co-expression under both promoters fully rescued the mutant phenotype (Fig. 6g). Given that the 3-kb regulatory region may lack distal elements required for endogenous *dmsr-2* expression in LUA, these findings are consistent with DMSR-2 functioning in LUA and additional neuronal populations to sustain the persistent behavioral state, although differences in receptor expression levels cannot be excluded.

To determine whether DMSR-2 also contributes to persistent circuit activity, we examined calcium dynamics following RnB stimulation. Similar to *nlp-13* mutants, *dmsr-2* mutants exhibited markedly reduced prolonged LUA calcium activity, whereas stimulus-evoked responses in R2B and R9B remained intact (Supplementary Fig. 9). Thus, disruption of either NLP-13 or DMSR-2 compromises sustained downstream activity without substantially affecting sensory-network responses. Together, these findings support a model in which LUA-derived NLP-13 engages DMSR-2-dependent circuit signaling to maintain prolonged neural activity and behavioral persistence.

## Discussion

Persistent behavioral states allow transient sensory experiences to influence subsequent actions after the initiating stimulus has disappeared. Here, we show that brief physical contact with a hermaphrodite induces a local search state in *C. elegans* males that persists for several minutes. This state is characterized by increased reversals and altered behavioral patterns, resulting in restricted locomotor coverage. By concentrating exploration near the site of a recent encounter, this behavioral strategy may increase the probability of relocating a mate after contact is temporarily lost. Importantly, entry into the local search state depends on recognition of a suitable mate rather than physical contact alone. Neither isolated cuticular material nor contact with a mechanically comparable but unsuitable nematode was sufficient to induce the state. These findings are consistent with our previous model in which males integrate cuticular cues and body stiffness to recognize compatible mates ^21^. Thus, local search appears to be gated by the convergence of multiple sensory features, preventing prolonged behavioral responses to irrelevant physical encounters. Our findings further identify glutamatergic and neuropeptidergic mechanisms that differentially contribute to sensory-network activation, prolonged LUA activity, and behavioral persistence (Fig. 7).

Transient RnB activation was sufficient to reproduce the persistent local search state induced by mate contact, identifying RnBs as the principal sensory population capable of initiating this response. In contrast, RnA activation alone failed to evoke comparable persistence. Increasing RnB stimulation progressively enhanced the magnitude and duration of local search, indicating that behavioral persistence is graded by the extent of sensory activation ^26^. Such stimulus-dependent recruitment may allow males to distinguish sustained interactions with suitable mates from incidental contact and engage persistent search when sensory activity exceeds an effective threshold. Comparable computational strategies operate across animal systems. In *Drosophila*, gustatory and olfactory pheromone pathways converge onto P1 neurons, where balanced excitation and inhibition regulate courtship initiation ^46^. In larval zebrafish, sensory evidence accumulates across distributed neural populations to regulate behavioral decisions ^47, 48^, while distinct sensory modalities coordinate social interactions across different spatial scales ^49^. Our findings suggest that distributed RnB activity similarly regulates state entry based on sensory input strength, although the precise integration mechanism remains to be determined.

Our results suggest that glutamatergic and neuropeptidergic signaling make distinct but potentially overlapping contributions to persistent behavioral states. Glutamatergic transmission facilitates neuronal responses within the interconnected RnB sensory network and contributes to downstream LUA activity (Fig. 7). In contrast, disruption of NLP-13 or its receptor DMSR-2 reduces prolonged LUA calcium activity and behavioral persistence while largely preserving the initial stimulus-evoked sensory responses. These observations are consistent with a model in which glutamatergic signaling contributes to establishing the local search response, whereas NLP-13/DMSR-2 signaling supports its persistence. However, the two signaling pathways may act across overlapping temporal phases, and their respective contributions to state initiation and maintenance remain to be further resolved. A related separation between initiation and maintenance mechanisms has been described in *Drosophila*, where P1 neurons initiate persistent courtship- and aggression-related states, while downstream integrative populations, including pCd neurons, contribute to behavioral persistence ^3, 6, 10, 50^. Persistent states can also involve prolonged recruitment of distributed neuronal populations, as observed during brain-wide state transitions in zebrafish ^51^. Together, these findings suggest that persistent behavioral states may arise from coordinated sensory-network recruitment and mechanisms that prolong downstream circuit activity, without necessarily requiring temporally exclusive signaling pathways.

Neuromodulatory signaling provides a potential mechanism for extending circuit activity beyond transient sensory input. In *C. elegans*, serotonergic and peptidergic pathways regulate transitions between roaming and dwelling states ^8^, while long-lasting neuropeptide signaling modulates female sexual drive in *Drosophila* ^52^. Neuropeptides are particularly suited to sustaining behavioral states because their actions can extend beyond individual synaptic events and influence distributed circuit components ^13, 14, 15, 16^. Interestingly, recent work in mammals suggests a related mechanism in hypothalamic circuits controlling persistent internal states ^53^. In mouse VMH^Esr^^1^ neurons, a predominantly glutamatergic population involved in aggression, disruption of oxytocin and vasopressin receptor signaling reduces persistent population activity and underlying attractor dynamics while only modestly affecting overall neuronal responsiveness. This resemblance suggests that coordination between fast excitatory transmission and slower neuropeptide signaling may represent a broadly employed strategy for sustaining neural activity. Our findings provide a cellularly defined example in which glutamatergic and neuropeptidergic signaling make complementary contributions to generating and sustaining persistent behavioral responses.

The cellular architecture through which NLP-13/DMSR-2 signaling maintains LUA activity remains unresolved. LUA-specific expression of NLP-13 rescues behavioral persistence, whereas expression in neighboring PHA neurons does not, indicating that the cellular source of the neuropeptide is functionally important. In contrast, rescuing the *dmsr-2* mutant phenotype required co-expression under a LUA-specific promoter and a broader 3-kb endogenous regulatory region (Fig. 6g). Although differences in receptor expression levels may contribute to this requirement, the combinatorial rescue is consistent with DMSR-2 acting in multiple circuit populations to sustain behavioral persistence. One possibility is that LUA-derived NLP-13 engages both local feedback through DMSR-2 in LUA and broader paracrine signaling to DMSR-2-expressing partner neurons. These additional neurons may provide recurrent excitation to LUA or modulate downstream premotor circuits, thereby stabilizing the persistent behavioral state. Alternatively, NLP-13/DMSR-2 signaling may prolong LUA activity by modulating intrinsic neuronal excitability rather than through recurrent circuit excitation. Identifying the relevant DMSR-2-expressing neurons and their functional interactions with LUA will be essential to distinguish these possibilities and define the circuit architecture underlying peptidergic maintenance.

Together, our findings illustrate how glutamatergic and neuropeptidergic signaling cooperate to transform brief sensory encounters into prolonged neuronal responses and persistent behavioral states. Glutamatergic signaling contributes to sensory-network recruitment and downstream LUA responses, whereas NLP-13/DMSR-2 signaling supports prolonged circuit activity and behavioral persistence. Although these pathways may operate across overlapping temporal phases, their complementary contributions provide a framework for understanding how neural circuits convert transient sensory experiences into sustained behavioral responses.

## Methods

### *C. elegans* strains and maintenance

*C. elegans* strains were maintained on nematode growth medium (NGM) plates seeded with *Escherichia coli* OP50 at 20 °C unless otherwise indicated, following standard culture procedures ^54^. A complete list of strains used in this study is provided in Supplementary Table 1.

### Molecular cloning

All expression constructs were generated by PCR amplification of the indicated promoter, coding, or genomic sequences and assembled into the pPD95.75 vector backbone using Gibson assembly. A complete list of primers is provided in Supplementary Table 2.

For preparation of genomic DNA templates, ten adult hermaphrodites were transferred into PCR tubes containing 10 μL freshly prepared lysis buffer consisting of 50 mM KCl, 10 mM Tris-HCl (pH 8.3), 2.5 mM MgCl_2_, 0.45% Tween-20, 0.45% NP-40, 0.01% gelatin, and 1 mg/mL proteinase K. Animals were lysed at 60°C for 1 h, followed by incubation at 95°C for 15 min to inactivate proteinase K. The resulting lysates were used directly as genomic DNA templates.

For preparation of cDNA templates, total RNA was extracted using the GENEzol™ TriRNA Pure Kit (Geneaid, Cat. No. GZX050). First-strand cDNA was synthesized using the SuperScript® III First-Strand Synthesis System for RT-PCR (Invitrogen, Cat. No. 18080-051) according to the manufacturer’s instructions. Promoter regions used for cell-specific expression and their corresponding lengths are listed in Supplementary Table 2.

### Tissue-specific RNAi

Tissue-specific RNAi was performed by expressing complementary sense and antisense fragments of the target gene in the indicated neurons. A fragment of the *eat-4* coding sequence was amplified and cloned in both orientations downstream of either an RnB-specific (*pkd-2*) or LUA-specific promoter (*C39H4.2*). The corresponding sense and antisense constructs were co-injected into the gonads of young adult hermaphrodites together with a fluorescent co-injection marker. Transgenic progeny carrying the co-injected constructs were identified using the fluorescent co-injection marker and examined in behavioral assays.

### Optogenetics and behavioral tracking

For optogenetic experiments, L4 males were transferred 24 h before the assay to NGM plates seeded with 100 μL *E. coli* OP50 supplemented with 2 mM all-trans-retinal (ATR; MedChemExpress, Cat. No. HY-W004500) and maintained in darkness until testing. Control animals were prepared in parallel without ATR.

Behavioral assays were performed using the WormLab tracking system (MBF Bioscience). Individual day-1 adult males were transferred to NGM assay plates seeded with 20 μL OP50 and allowed to move freely while being recorded. Chrimson-expressing neurons were activated with 595-nm red light at an irradiance of 0.428 mW/mm². Unless otherwise indicated, stimulation consisted of 500-ms light pulses separated by a 1-s interval. The number of pulses used in each experiment is indicated in the corresponding figure or schematic. Behavior was recorded for 10 min, encompassing baseline, stimulation, and post-stimulation periods. Animal position, locomotor trajectories, reversals, turns, and related locomotor parameters were extracted using WormLab software.

### Contact and behavioral tracking

Contact-induced local search was examined using individual day-1 adult males. Each male was transferred to an NGM assay plate and allowed to move freely before presentation of a potential mate. An immobilized *unc-13* mutant hermaphrodite was positioned immediately adjacent to the moving male using an eyelash. Because males could make brief or intermittent contacts before establishing sustained interaction, successful contact was operationally defined as 30 s of continuous physical contact with the hermaphrodite. Brief contacts occurring before this criterion was met were retained as part of the natural interaction sequence and were not considered the onset of the defined 30-s contact period. After 30 s of continuous contact, the hermaphrodite was removed and male locomotion was recorded during the post-contact period. For the mock-treated no-contact condition, an *unc-13* hermaphrodite was placed near the male without permitting physical contact. Where indicated, males were tested using *daf-22* mutant hermaphrodites, purified hermaphrodite cuticles, or *Pristionchus pacificus* as contact stimuli.

Behavior was recorded for 10 min using the WormLab tracking system (MBF Bioscience) or a CCD camera (Touptek XCAMTOP4K8MPB). The centroid position of each animal was extracted throughout the recording and used to reconstruct locomotor trajectories. Reversals, turns, instantaneous speed, and other locomotor parameters were extracted using WormLab.

### CASTLE behavioral classification

Individual 10-min open-field recordings (acquired at 7.5 frames/s), or datasets comprising 15 recordings as indicated, were analyzed using the CASTLE behavioral-classification pipeline. Regions of interest containing individual animals were segmented and tracked throughout each recording. Background subtraction was performed to remove residual locomotion trails, and animal orientation was computationally normalized before extraction of focused visual latent representations. These latent features were analyzed together with conventional kinematic measurements, including x-y position, instantaneous speed, region-of-interest area, and angular speed.

Behavioral features were processed using a two-stage dimensionality-reduction procedure based on Uniform Manifold Approximation and Projection (UMAP). In the first stage, latent representations were reduced using UMAP with *n_neighbors* = 500 and *n_components* = 12, followed by a second stage with *n_neighbors* = 50 and *n_components* = 2. Behavioral observations were then assigned to the nearest existing cluster as reported ^27^. The resulting behavioral clusters were assigned to global search, local search, looping, and curling classes based on manual confirmation from blind tests.

### Quantification of locomotor dynamics and heatmap visualization

Locomotor trajectories were obtained using WormLab by tracking the center position of each animal throughout the recording. To quantify changes in net locomotor displacement over time, trajectories were divided into consecutive 10-s bins. For recordings acquired at 7.5 frames/s, each bin corresponded to 75 frames. Net displacement within each bin was calculated as the Euclidean distance between the animal’s position at the beginning and end of the interval, thereby minimizing the contribution of small frame-to-frame tracking fluctuations.

For each animal, the pre-stimulation period was used to establish an individual baseline for spatial exploration. The mean displacement across the pre-stimulation analysis window (0-275 s) was calculated, and displacement in each 10-s bin was expressed as a ratio relative to this baseline (bin displacement/pre-stimulation mean displacement). A ratio of 1 therefore represents the animal’s own baseline level of exploration, whereas values below 1 indicate reduced spatial exploration. For dynamic heatmaps, these baseline-normalized ratios were plotted for individual animals across time, with animals displayed as separate rows and time bins arranged chronologically. The stimulation or contact period itself was excluded from baseline and post-stimulation quantification. For analyses comparing pre- and post-event dynamics, equivalent 275-s windows before and after the event were used.

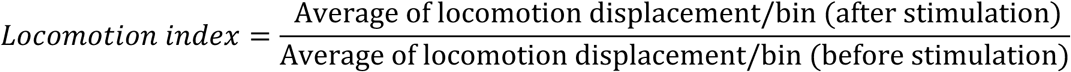

### Quantification of behavioral recovery dynamics

Time to baseline recovery was determined independently for each animal from the 10-s binned displacement trace. The mean displacement during the pre-stimulation analysis window was defined as that animal’s baseline. Beginning after the end of stimulation or mate contact, behavioral recovery was defined as the first time at which displacement returned to or exceeded the individual baseline for two consecutive 10-s bins. The recovery time was assigned to the beginning of the first bin in this two-bin sequence, thereby requiring recovery to be sustained for at least 20 s and reducing the likelihood that a single transient excursion above baseline would be scored as recovery. Animals that did not meet this criterion during the post-stimulation observation period were classified as non-recovered and censored at the end of the observation window for time-to-recovery analysis. Because this recovery criterion was applied to all animals regardless of the magnitude of their initial behavioral response, time to baseline recovery was interpreted as an operational measure of locomotor recovery rather than a direct measurement of persistent-state duration in every animal.

Time-to-recovery distributions were visualized using Kaplan–Meier curves, in which the survival function represents the fraction of animals that had not yet met the predefined baseline-recovery criterion. Animals that failed to recover within the observation window were treated as right-censored at the final time point (275 s in the post-stimulation). Differences between experimental groups were assessed using the log-rank (Mantel-Cox) test. Recovery proportions were additionally reported as the number and percentage of animals satisfying the recovery criterion within the observation period.

### Chemogenetic neuronal silencing

For chemogenetic silencing experiments, a 1 M histamine stock solution was prepared and adjusted to pH 6.3. Histamine (Sigma, Cat. No. H7250) was added to NGM agar to obtain a final concentration of 1 mM. L4 males expressing the histamine-gated chloride channel HisCl1 in the indicated neurons were transferred to histamine-containing NGM or regular NGM plates seeded with OP50 and maintained for 24 h before behavioral testing. Control animals carrying the same transgene were maintained under identical conditions in the presence or absence of histamine. For experiments combining chemogenetic silencing with optogenetic stimulation, animals were also supplied with 2 mM ATR during the 24-h conditioning period.

### Calcium imaging

Calcium imaging was performed in day-1 adult males expressing GCaMP8f under the control of either a *pkd-2*-associated promoter or a LUA-specific promoter in a *lite-1(ce314)* background. Animals were immobilized by microbeads (Polyscience, Cat. No. 00876-15) and imaged using a ZEISS Axio Imager M2 microscope equipped with an Axiocam 710 monochrome camera and a 63X objective.

GCaMP8f fluorescence images were acquired at 1-s intervals with an exposure time of 40 ms. For optogenetic stimulation, neurons were illuminated with 620-nm light for 500 ms. GCaMP8f fluorescence images were acquired continuously immediately after optogenetic stimulation. Animals received 20 pulses (500 ms/pulse) of optogenetic stimulation at 1-s intervals, or as indicated for each experiment. Recordings consisted of 240 frames (1 Hz), including 40 s of pre-stimulation baseline followed by stimulation and post-stimulation periods.

For fluorescence quantification, regions of interest (ROIs) were manually defined around the cell bodies of LUA or RnB neurons. An adjacent region of comparable size containing no detectable neuronal fluorescence was selected as the background ROI. Background fluorescence was subtracted from the neuronal ROI at each time point. ROIs were tracked throughout the recording to account for animal or neuronal movement. Fluorescence intensity was quantified using Fiji ^55^. Baseline fluorescence (*F_0_*) was calculated as the mean background-subtracted fluorescence intensity over the first 40 s acquired before optogenetic stimulation. Calcium responses were calculated as:

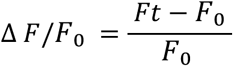

where *Ft* represents the fluorescence intensity at each time point and *F_0_* represents the mean pre-stimulation baseline fluorescence. Peak response amplitude and the persistence of post-stimulation calcium activity were quantified as indicated in the corresponding figure legends.

Fluorescence traces were analyzed without photobleaching correction. Although gradual fluorescence decay was observed during some recordings, fitting the pre-stimulation fluorescence decay and extrapolating this function across the recording introduced a systematic upward distortion of the post-stimulation signal. We therefore retained the raw background-subtracted fluorescence measurements and normalized them to the pre-stimulation F_0_ (0-40 s). Stimulus-associated calcium responses were quantified as the baseline-adjusted area under the ΔF/F_0_ curve from 40 to 70 s. For each recording, a linear reference was constructed between the ΔF/F_0_ values at the beginning and end of this interval, and the area between the recorded trace and this reference was calculated. Prolonged post-stimulation LUA calcium activity was quantified as follows. Calcium-dependent fluorescence signals were normalized to the prestimulation baseline (F_0_) to obtain ΔF/F_0_ for each animal. To account for time-dependent fluorescence changes observed in controls, the mean normalized fluorescence of animals recorded without all-trans retinal (ATR-) was calculated at each frame and subtracted from the corresponding fluorescence value of each experimental animal. Prolonged fluorescence responses were quantified by averaging the corrected values over 120-240 s:

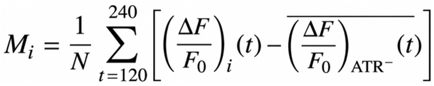

where (M_i_) represents the mean ATR-negative-control-corrected fluorescence response of animal (i) over the analysis interval, and (N) is the number of frames included in that interval.

### Calcium-response heatmap visualization

For heatmap visualization, calcium traces from individual animals were restricted to 30-240 s to focus the analysis on the pre-stimulation, stimulation, and post-stimulation periods. For each individual trace, fluorescence values within this interval were independently standardized by z-score transformation. Specifically, the mean and standard deviation were calculated across 30-240 s for that animal, and the signal at each frame was expressed as the number of standard deviations above or below that trace-specific mean. Thus, z-score normalization was performed independently for each animal and was used only to visualize the temporal pattern and relative magnitude of calcium dynamics, rather than to compare absolute fluorescence amplitudes between animals. The resulting z-scored traces were displayed as heatmaps, with individual animals arranged as separate rows and time points shown chronologically from 30 to 240 s.

### Confocal and epifluorescent microscopy

Confocal images were acquired using an LSM 780 confocal microscope (Zeiss) and analyzed using ZEN 3.6 software (Zeiss). Epifluorescence images were acquired using a Zeiss Axio Imager M2 microscope equipped with an Axiocam 710 monochrome camera.

### Quantification of NLP-13::mKate fluorescence

To examine contact-dependent changes in NLP-13 reporter fluorescence, day-1 adult males expressing NLP-13::mKate in LUA neurons were assigned to control and contact-exposed groups. Males in the contact-exposed group were incubated with adult hermaphrodites for 2 h, whereas control males were maintained under otherwise identical conditions without hermaphrodites. Animals were subsequently immobilized using polystyrene microbeads and imaged with a Zeiss Axio Imager M2 microscope equipped with an Axiocam 710 monochrome camera. All animals within an experiment were imaged using identical objective, illumination intensity, exposure time, camera-gain, and acquisition settings.

NLP-13::mKate fluorescence was quantified using Zen. A region of interest was manually drawn around the LUA cell body, and an adjacent region of comparable size lacking detectable neuronal fluorescence was used to estimate background intensity. Background-corrected fluorescence was calculated by subtracting the mean intensity of the background region from the mean fluorescence intensity of the LUA region. The same ROI-selection and background-subtraction procedures were applied to all animals.

### Statistical analyses

Statistical analyses were performed using GraphPad Prism version 10.4.1. The statistical test used for each experiment is specified in the corresponding figure legend. Comparisons between two groups were performed using unpaired or paired two-tailed Student’s t-tests where appropriate. Comparisons among three or more groups with a single experimental factor were performed using one-way ANOVA followed by Bonferroni’s multiple-comparisons test. Experiments involving multiple groups or experimental factors were analyzed using two-way ANOVA, followed by Bonferroni’s multiple comparisons test. For analyses of time to baseline displacement recovery, time-to-recovery distributions were plotted as Kaplan-Meier curves and compared using the log-rank (Mantel-Cox) test. Animals that did not recover within the observation period were treated as right-censored. Individual data points represent independent animals unless otherwise specified. Sample sizes and the number of independent biological replicates are reported in the corresponding figure legends. Statistical significance was defined as *p*< 0.05. Statistical analyses of locomotor and calcium-imaging measurements were performed using individual animals as independent biological units, with repeated measurements from each animal summarized before between-group comparisons.

### Declaration of Generative AI and AI-assisted Technologies in the Writing Process

During the preparation of this work, the authors used ChatGPT to improve the language, grammar, and readability of the manuscript. After using this tool/service, the authors reviewed and edited the content as needed and take full responsibility for the content and integrity of the final publication.

## Data availability

The data supporting the findings of this study are available within the article and its Supplementary Information. Strains and plasmids generated in this study are available from the corresponding author upon reasonable request.

## Acknowledgments

We thank the members of the Chen and Sternberg labs, Dr. Han Wang, Dr. Chun-Liang Pan, and Dr. Vladislav Susoy for their insightful discussions, and Clementine Chou for her efforts in data processing. We also thank Chun-Liang Pan for sharing reagents. Some strains were provided by the *C. elegans* Genetics Center, funded by the NIH Office of Research Infrastructure Programs (P40 OD010440). We are grateful for the technical assistance of Technology Commons, College of Life Science, National Taiwan University, with confocal microscopy (CLSM). We thank the *C. elegans* Core Facility of the National Core Facility for Biopharmaceuticals, National Science and Technology Council (NSTC) in Taiwan. This study was funded by the National Science and Technology Council (NSTC) in Taiwan, to C.-H.C. (NSTC 109-2636-B-002-016; NSTC 110-2636-B-002-013; NSTC 111-2636-B-002-027; NSTC 112-2636-B-002-007; NSTC 113-2636-B-002-003; NSTC 114-2311-B-002-024; NSTC 115-2311-B-002-016).

## Author contribution

Conceptualization, Y.-H. C., Y.-C. L., T.-Y. C., K.-A. Y., and C.-H. C.; methodology, Y.-H. C., Y.-C. L., T.-Y. C., K.-A. Y., and J.-B. Y. ; Investigation, Y.-H. C., Y.-C. L., T.-Y. C., K.-A. Y., W.-E. M., Y.-S. L. and C. W.; funding acquisition, C.-H. C.; Writing-original draft, Y.-H. C., C.-H.C.; Writing-review and editing, Y.-H. C., C.-H. C.

