## Supplementary Information for "A Glutamate-Neuropeptide Circuit Converts Transient Mate Contact into a Persistent Local Search State in *Caenorhabditis elegans* Males"

9 Supplementary Figures

2 Supplementary Tables

6 Supplementary Videos

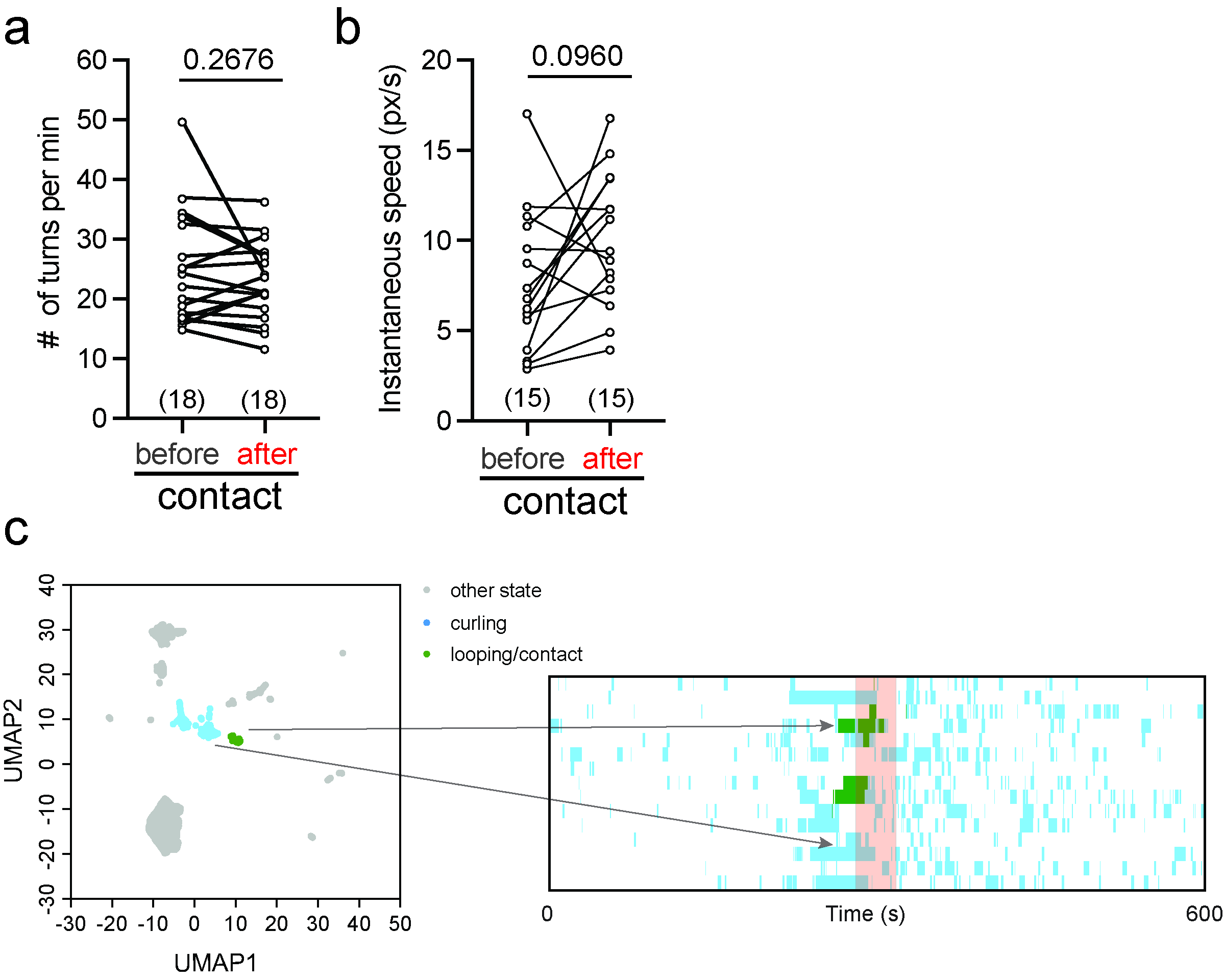

**Supplementary Fig. 1 Mate contact alters behavioral-state organization without significantly changing turning frequency or instantaneous speed.**

a, Number of turns per minute in males before and after 30-s contact with a mate (*n* = 18). b, Instantaneous locomotion speed before and after mate contact (*n* = 15). Lines connect measurements from the same animal. Statistical significance was assessed using two-tailed paired Student’s *t*-tests; exact *P* values are shown. c, UMAP-based behavioral classification following mate contact, showing curling (blue), looping/contact (green), and other behavioral states (grey). The corresponding temporal behavioral-state sequences during the 600-s assay are shown at right. Lines indicate representative behavioral epochs mapped between the UMAP space and the temporal classification.

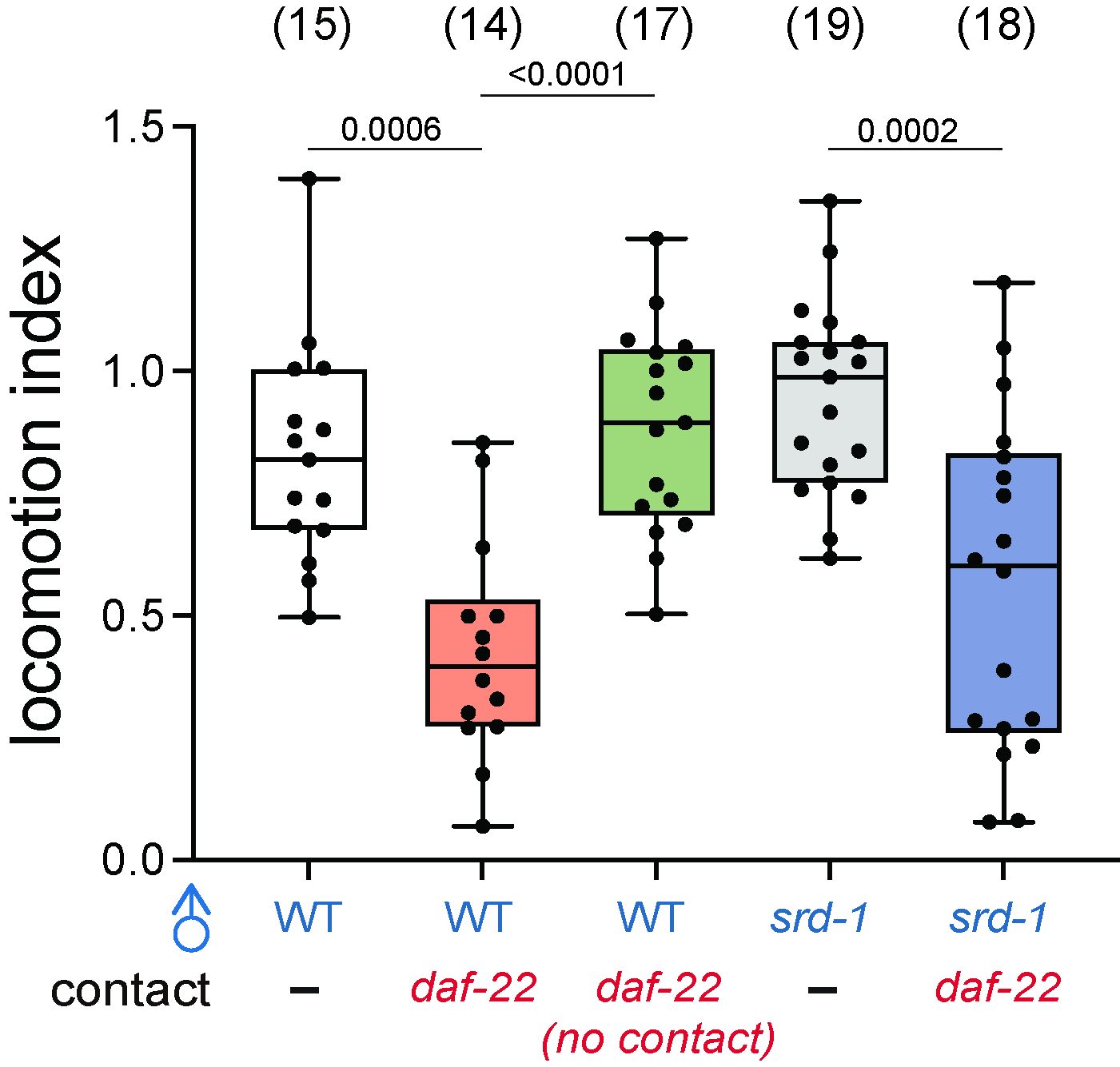

**Supplementary Fig. 2 Ascaroside and SRD-1-dependent pheromone signaling are dispensable for contact-induced local search.**

Locomotion indices of wild-type and *srd-1* mutant males in the absence of a mate (-) or following exposure to *daf-22* mutant hermaphrodites (with or without physical contact). *daf-22* mutant hermaphrodites were also placed away from males as another control. Boxes indicate the median and 25th-75th percentiles, whiskers indicate the minimum and maximum values, and dots represent individual animals. Sample sizes are indicated above each group. Statistical significance was assessed by one-way ANOVA followed by Bonferroni correction for multiple comparisons; exact *P* values are shown.

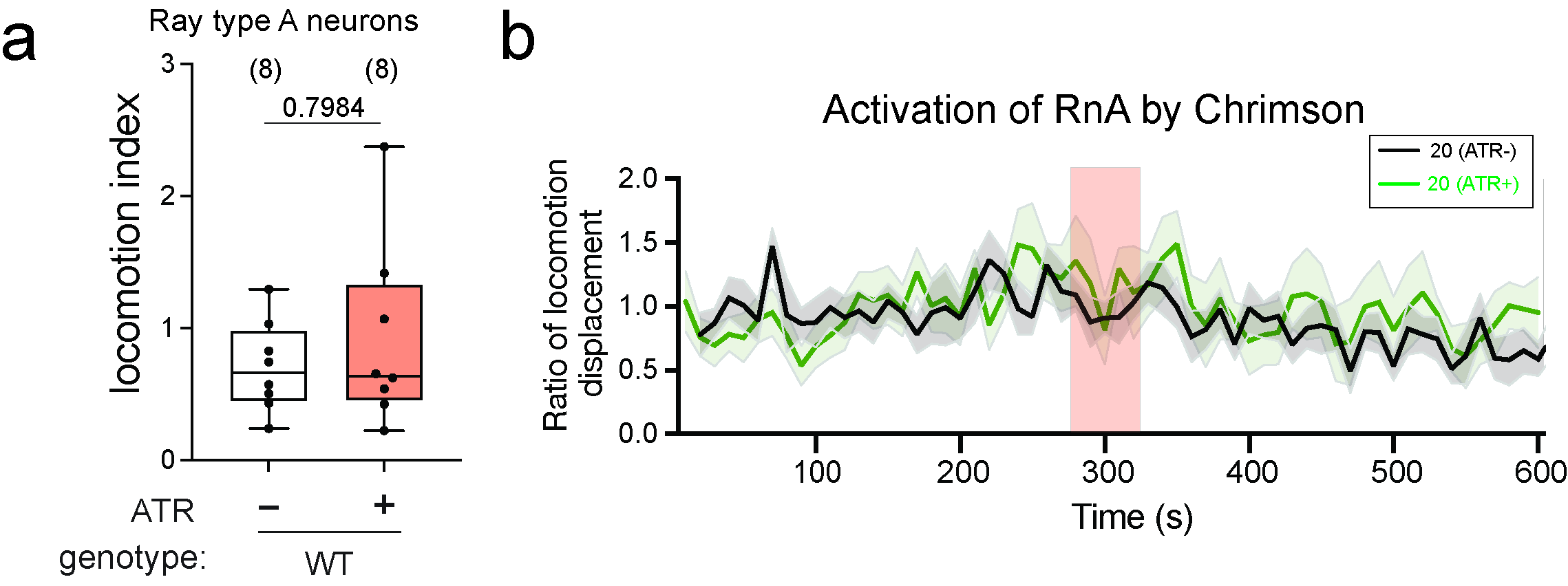

**Supplementary Fig. 3 Optogenetic activation of ray type A neurons is insufficient to induce persistent local search.**

a, Locomotion indices of males expressing Chrimson in ray type A neurons (RnAs) under ATR-negative and ATR-positive conditions following 20-pulse optogenetic stimulation (*n* = 8 per condition). Boxes indicate the median and 25th-75th percentiles, whiskers indicate the minimum and maximum values, and dots represent individual animals. Statistical significance was assessed by the two-tailed unpaired Student's t-test; the exact *P* value is shown. b, Time course of the locomotion displacement ratio following 20-pulse optogenetic stimulation of RnAs in ATR-negative and ATR-positive animals. Lines indicate mean responses and shaded regions indicate ± SEM.

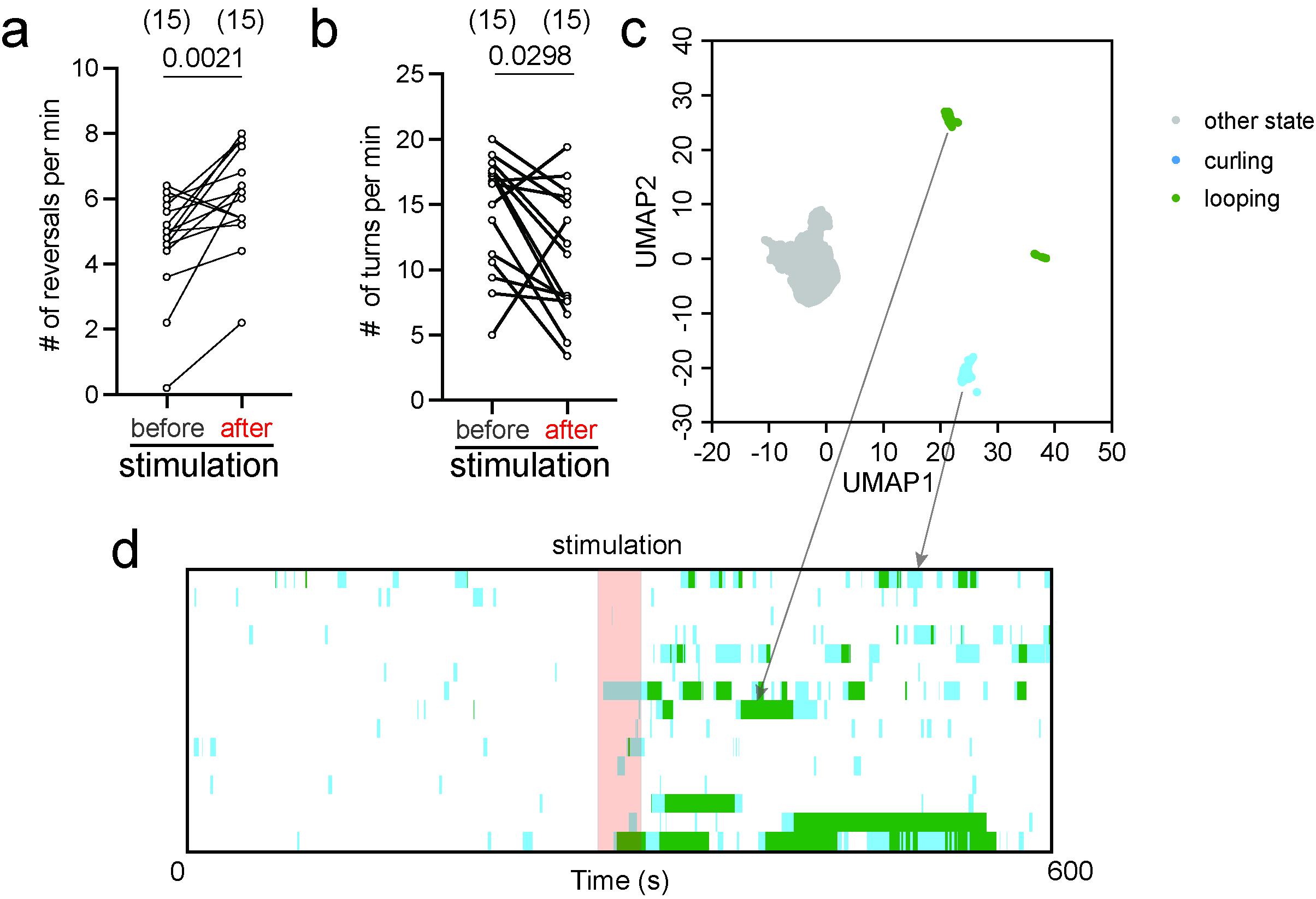

**Supplementary Fig. 4 RnB activation increases reversals and reorganizes post-stimulation behavioral states.**

a, b, Number of reversals (a) and turns (b) per minute before and after optogenetic stimulation of RnB neurons (*n* = 15). Lines connect measurements from the same animal. Statistical significance was assessed using two-tailed paired Student’s *t*-tests; exact *P* values are shown. c, UMAP-based classification of behavioral states following RnB activation, showing curling (blue), looping (green), and other behavioral states (grey). d, Temporal behavioral-state sequences of individual animals during the 600-s assay. The shaded region denotes the stimulation period, and lines connect representative UMAP clusters with corresponding behavioral epochs.

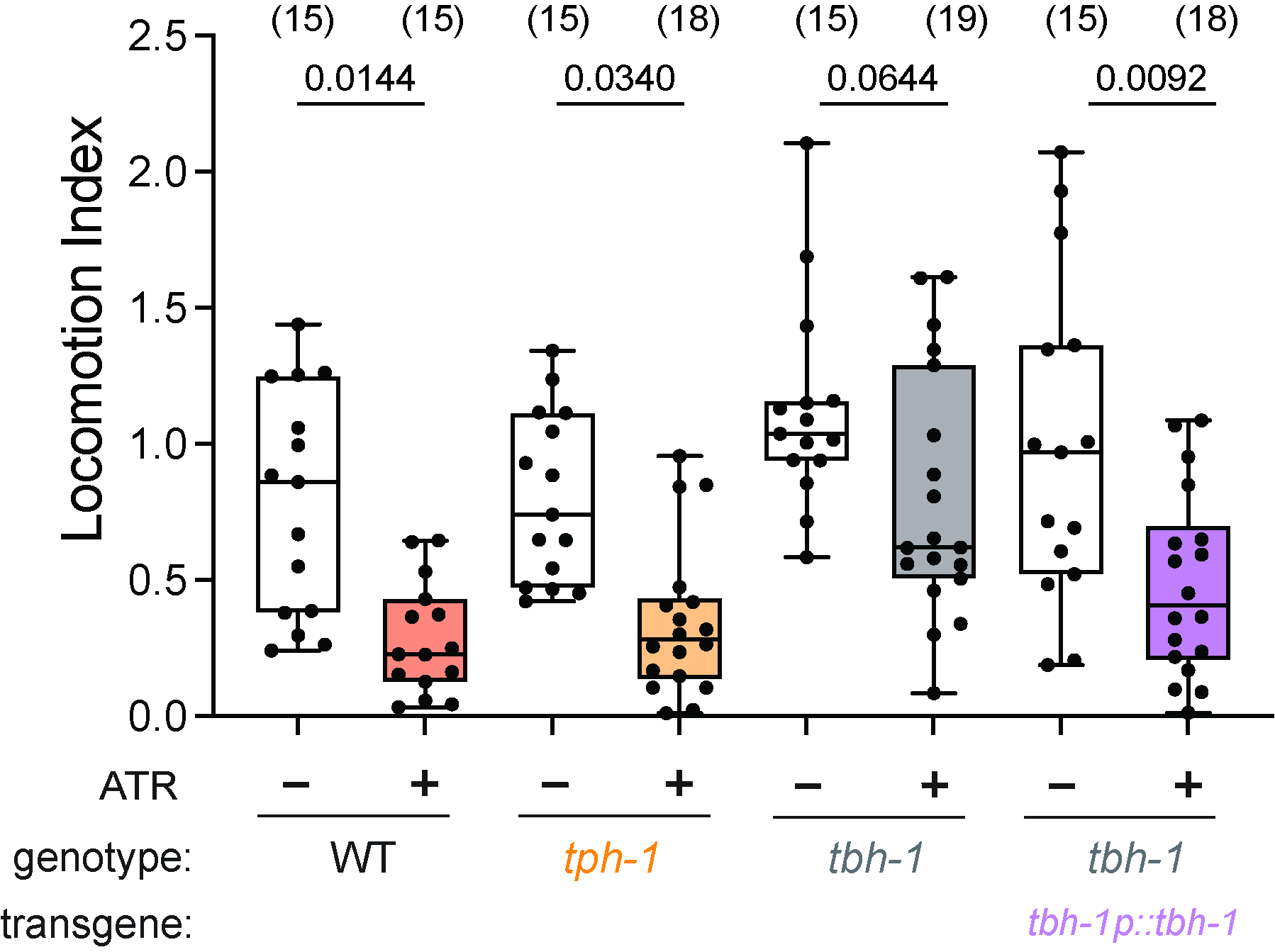

**Supplementary Fig. 5 Octopaminergic signaling modulates RnB-induced local search.**

Locomotion indices following RnB optogenetic stimulation in wild-type, *tph-1*, and *tbh-1* mutant males, and in *tbh-1* mutants carrying a *tbh-1p::tbh-1* rescue transgene. ATR-negative and ATR-positive conditions are shown. Boxes indicate the median and 25th-75th percentiles, whiskers indicate the minimum and maximum values, and dots represent individual animals. Sample sizes are indicated above each group. Statistical significance was assessed by two-way ANOVA followed by Bonferroni correction for multiple comparisons; exact *P* values are shown.

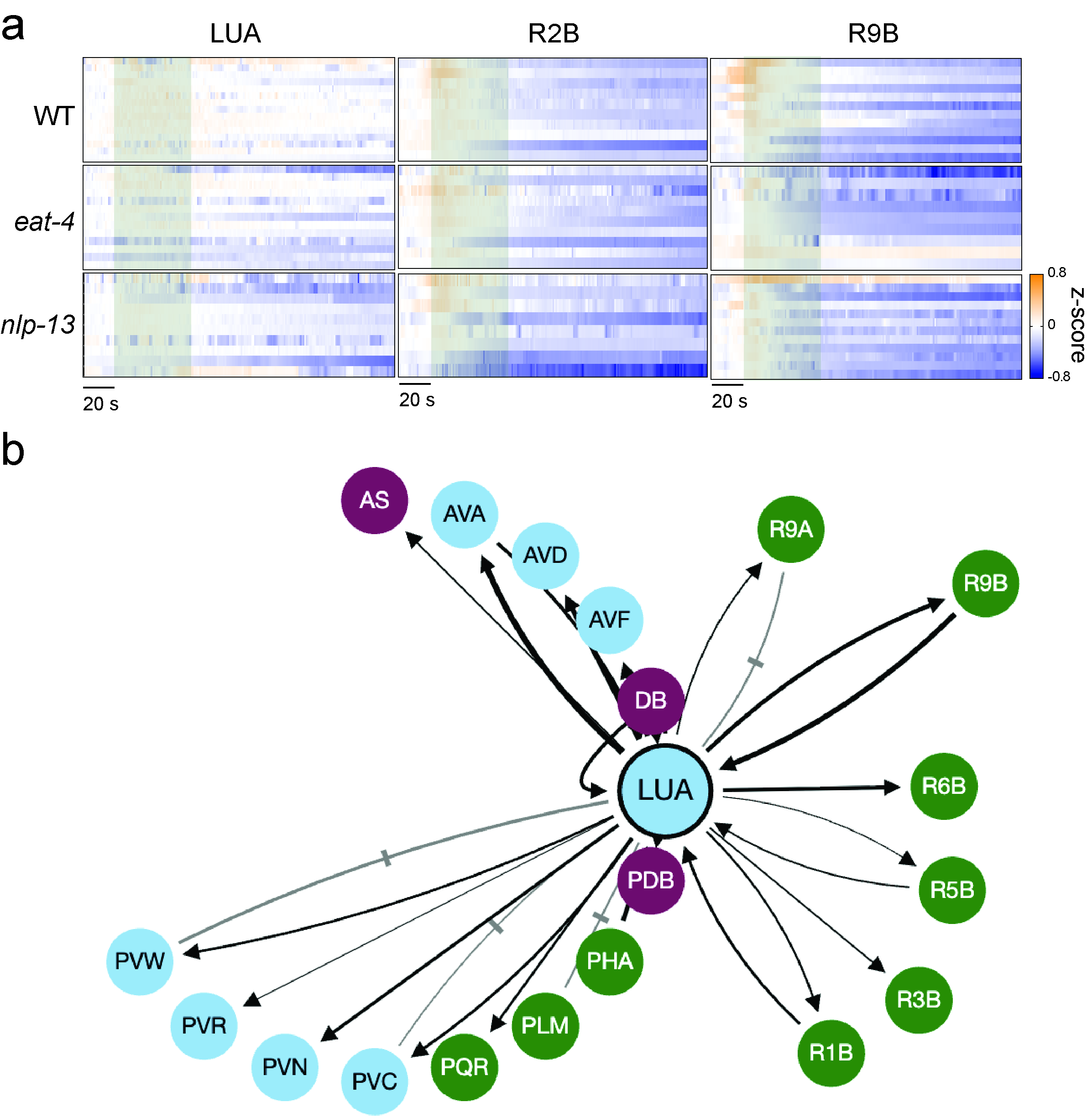

**Supplementary Fig. 6 Calcium-response dynamics and anatomical connectivity of the RnB-LUA circuit.**

a, Heatmaps of z-score-normalized calcium activity in LUA, R2B, and R9B neurons from wild-type, *eat-4*, and *nlp-13* males during RnB optogenetic stimulation. Individual animals are shown as separate rows and time points are arranged chronologically. b, Schematic of annotated synaptic connectivity between LUA and connected neurons, including multiple male-specific ray neurons. Arrows indicate the direction of chemical synaptic connectivity, grey lines with perpendicular bars indicate electrical synapses/gap junctions. Line thickness reflects relative connection strength where indicated. Neurons are grouped by neuronal class as shown in the schematic. Connectivity was reconstructed from the published *C. elegans* connectome dataset.

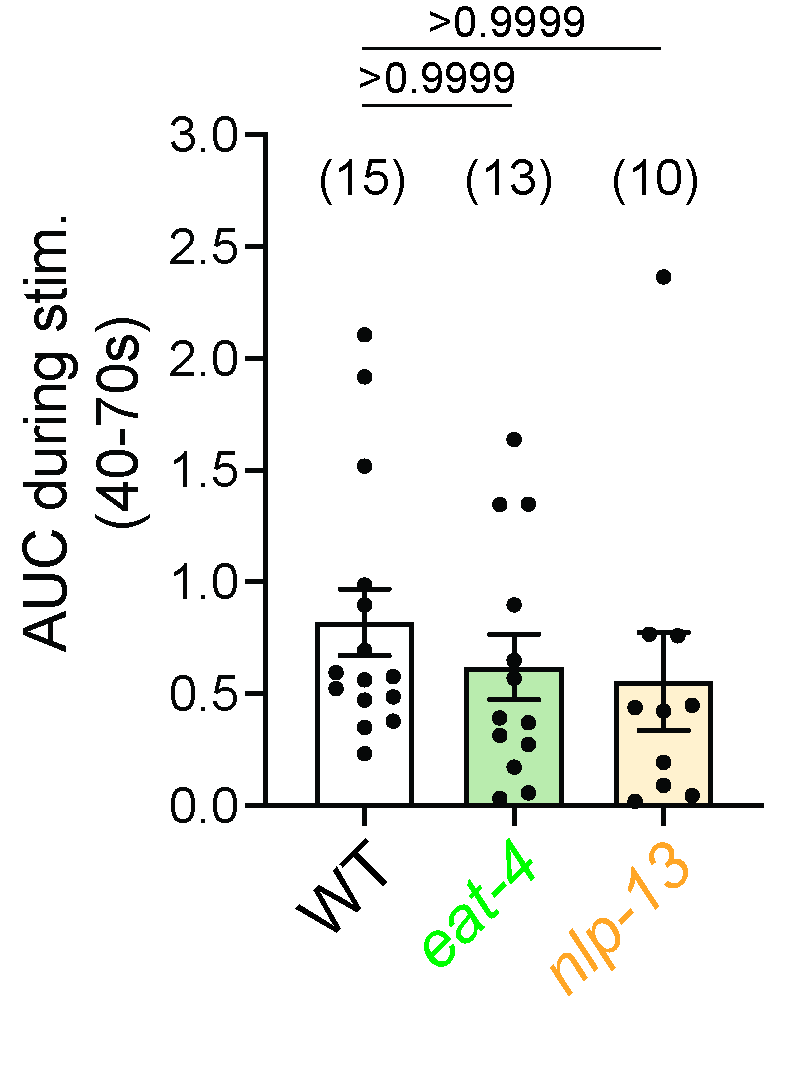

**Supplementary Fig. 7 *eat-4* and *nlp-13* do not significantly alter the stimulation-associated LUA calcium response.**

Area under the curve (AUC) of LUA calcium responses during the stimulation-associated interval (40-70 s) in wild-type, *eat-4*, and *nlp-13* males. AUC was calculated relative to the local linear baseline as described in the Methods. Data are mean ± SEM with individual animals shown. Statistical significance was assessed by one-way ANOVA followed by Bonferroni correction for multiple comparisons; exact *P* values are shown.

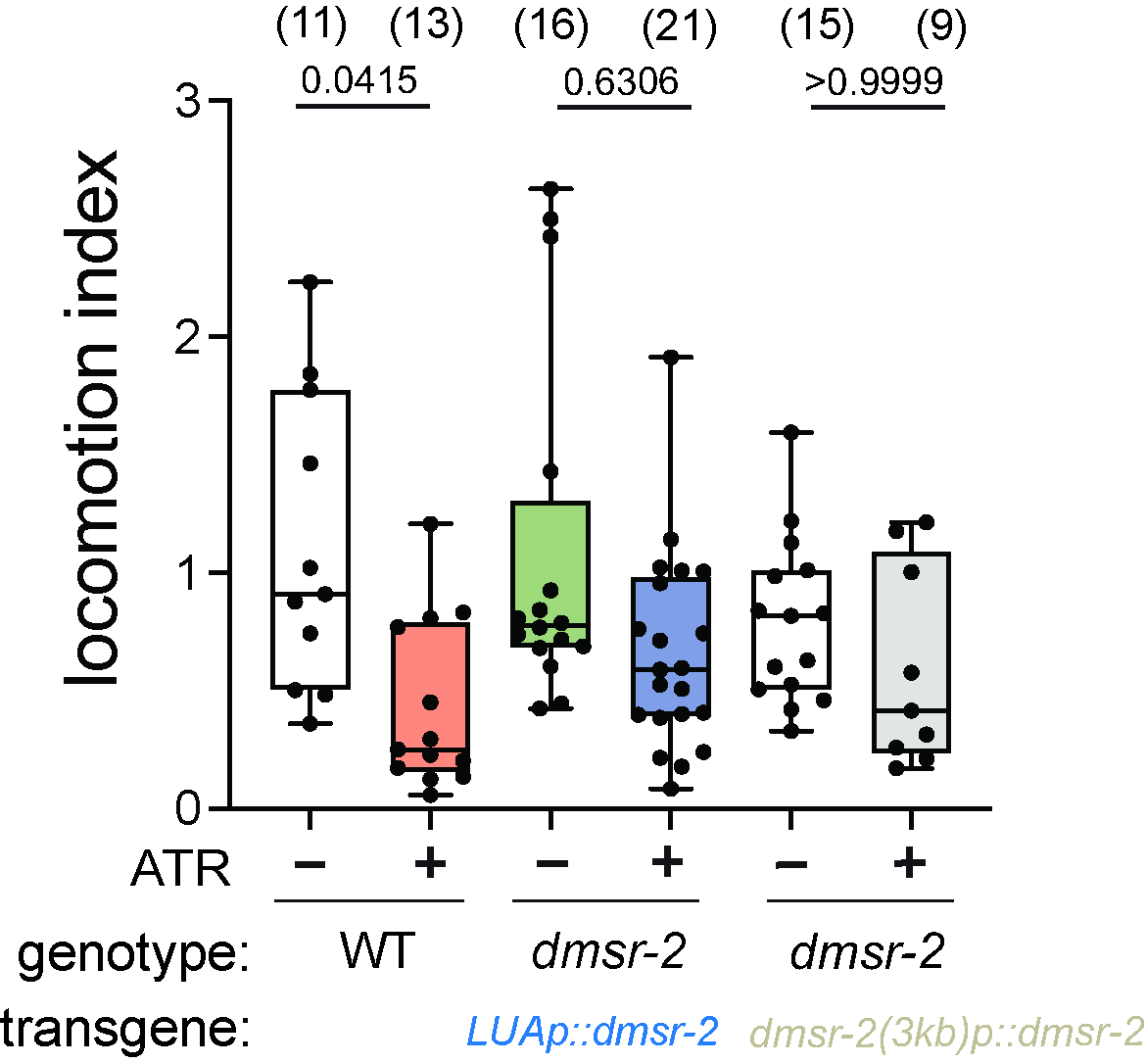

**Supplementary Fig. 8 Expression of DMSR-2 under individual promoters fails to rescue persistent behavioral responses in *dmsr-2* mutants.**

Locomotion indices of wild-type (WT) males and *dmsr-2* mutant males expressing *dmsr-2* under either an LUA-specific promoter (*LUAp::dmsr-2*) or a 3-kb endogenous upstream regulatory region (*dmsr-2(3kb)p::dmsr-2*), measured with or without all-trans retinal (ATR) following optogenetic stimulation of RnB neurons. ATR-dependent stimulation significantly reduced the locomotion index in WT males, whereas neither transgenic expression condition restored a significant ATR-dependent behavioral response in *dmsr-2* mutants. Box plots indicate the median (center line), interquartile range (box), and minimum and maximum values (whiskers), with individual data points overlaid. Sample sizes are indicated above each group. Statistical significance was determined by two-way ANOVA followed by Bonferroni's multiple-comparisons test. Exact *P* values are indicated above the corresponding comparisons.

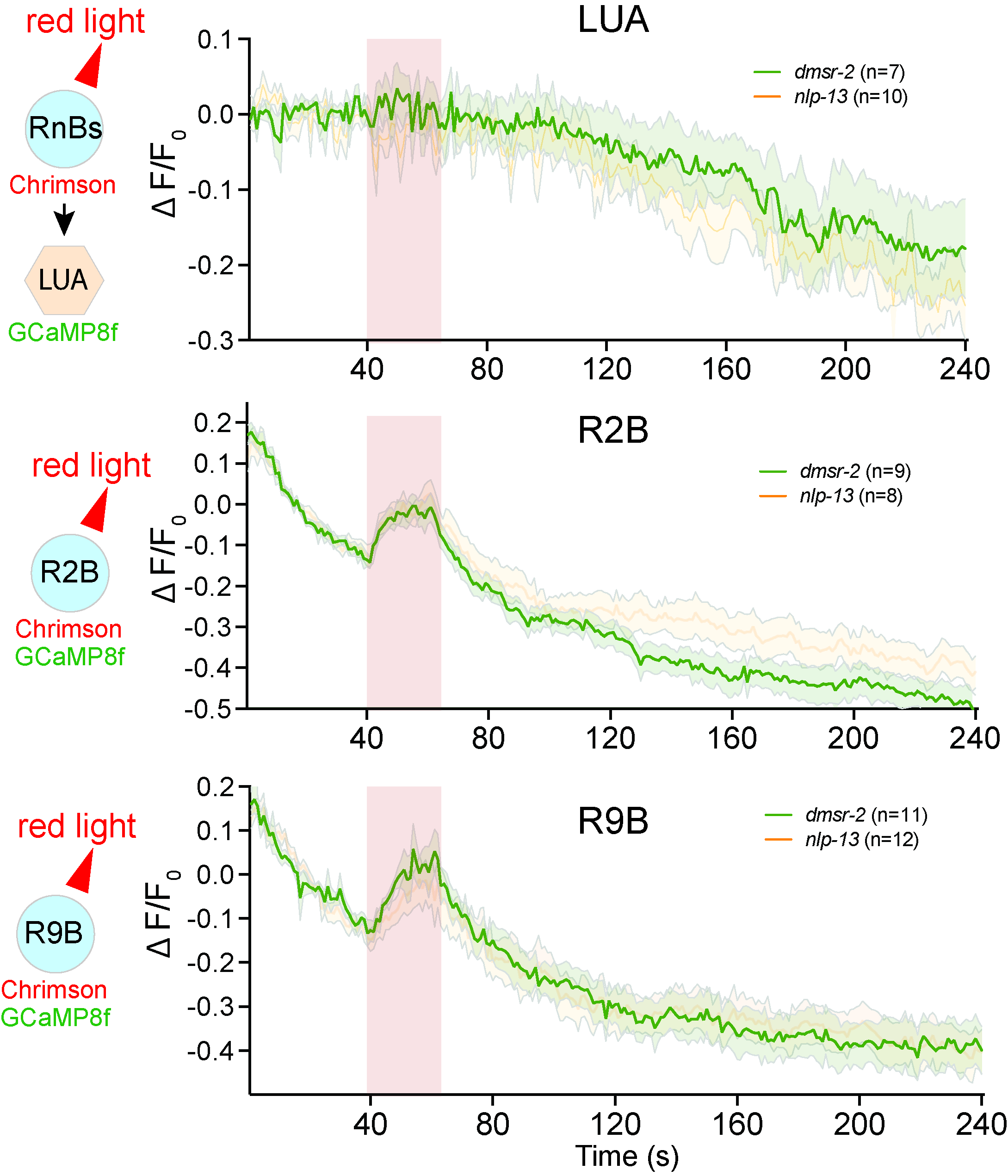

**Supplementary Fig. 9 *dmsr-2* mutant phenocopies *nlp-13* mutant phenotypes.**

Calcium responses of LUA, R2B, and R9B neurons during and after optogenetic activation of RnB neurons in *nlp-13* and *dmsr-2* mutant males. Traces show ΔF/F_0_ over time; sample sizes are indicated in the figure. Solid lines indicate mean responses and surrounding boundaries/shading indicate variability among animals. Traces of *nlp-13* were replotted from Fig. 5 for comparison.

**Supplementary Tables**

**Supplementary Table 1.** List of *C. elegans* strains used in this study.

**Supplementary Table 2.** List of primers used for molecular cloning in this study.

**Supplementary Videos**

**Supplementary Video 1.** Representative video showing mate-induced persistent local search behavior in a wild-type male following contact with an *unc-13* mutant hermaphrodite.

**Supplementary Video 2.** Representative video showing male behavioral responses to optogenetic activation of RnA neurons.

**Supplementary Video 3.** Representative video showing male behavioral responses to optogenetic activation of RnB neurons.

**Supplementary Video 4.** Representative video showing persistent local search behavior in a wild-type male following optogenetic activation of RnB neurons.

**Supplementary Video 5.** Representative video showing behavioral responses in an *eat-4* mutant male following optogenetic activation of RnB neurons.

**Supplementary Video 6.** Representative video showing behavioral responses in an *nlp-13* mutant male following optogenetic activation of RnB neurons.
